# TREM1 disrupts myeloid bioenergetics and cognitive function in aging and Alzheimer’s disease models

**DOI:** 10.1101/2024.03.05.583562

**Authors:** Edward N. Wilson, Congcong Wang, Michelle S. Swarovski, Kristy A. Zera, Hannah E. Ennerfelt, Qian Wang, Aisling Chaney, Esha Gauba, Javier A. Ramos Benitez, Yann Le Guen, Paras S. Minhas, Maharshi Panchal, Yuting J. Tan, Eran Blacher, Chinyere A. Iweka, Haley Cropper, Poorva Jain, Qingkun Liu, Swapnil S. Mehta, Abigail J. Zuckerman, Matthew Xin, Jacob Umans, Jolie Huang, Aarooran S. Durairaj, Geidy E. Serrano, Thomas G. Beach, Michael D. Greicius, Michelle L. James, Marion S. Buckwalter, Melanie R. McReynolds, Joshua D. Rabinowitz, Katrin I. Andreasson

## Abstract

Human genetics implicate defective myeloid responses in the development of late onset, age-associated Alzheimer’s disease (AD). Aging is characterized by a decline in myeloid metabolism that triggers maladaptive, neurotoxic immune responses. TREM1 is an amplifier of pro-inflammatory myeloid responses, and here we find that *Trem1* deficiency prevents age-dependent changes in myeloid metabolism, inflammation, and hippocampal memory function. *Trem1* deficiency rescues age-associated declines in ribose-5P, a glycolytic intermediate and the precursor for purine, pyrimidine, and NAD^+^ biosynthesis. In vitro, *Trem1* deficient microglia are resistant to bioenergetic changes induced by amyloid-ß_42_ oligomers (Aß_42_), suggesting that Aß_42_ stimulation disrupts homeostatic microglial metabolism and immune function via TREM1. In the 5XFAD model of amyloid accumulation, *Trem1* haploinsufficiency prevents spatial memory loss, preserves homeostatic microglial morphology, and reduces neuritic dystrophy independent of amyloid accumulation or changes in the disease-associated microglial transcriptomic signature. In aging *APP^Swe^* mice, *Trem1* deficiency restores synaptic mitochondrial function and cerebral glucose uptake and prevents hippocampal memory decline. In post-mortem human brain, microglial TREM1 expression increases with clinical and neuropathological severity. Thus, TREM1-mediated disruption of myeloid metabolism, both in the periphery and brain, promotes cognitive decline in aging and amyloid accumulation, two major risk factors for AD development.

## MAIN

Aging is the major risk factor for Alzheimer’s disease (AD)^1,2^ and is characterized by the development of persistent maladaptive inflammation^3,4^. Genome-wide association studies (GWAS) have identified genetic loci that associate with increased risk of AD^5^, many of which contain genes expressed in myeloid cells including brain microglia and peripheral monocytes and macrophages. This preponderance of myeloid genes suggests a pathogenic role for disrupted innate immunity in AD more consequential than previously appreciated. Preclinical studies in mouse models of aging and AD pathology demonstrate that healthy microglial function is lost with advancing age and suggest that disease-modifying components of the innate immune response could be targeted to slow or halt disease progression. A well-studied myeloid receptor associated with AD is TREM2 (Triggering Receptor Expressed on Myeloid cells-2) where loss of function variants increase risk for AD^6–8^. TREM2 is an anti-inflammatory innate immune receptor that promotes Aβ clearance^9,10^ and microglial survival^11^. In models of amyloid accumulation, microglia lacking *Trem2* or harboring the R47H *Trem2* variant show deficient proliferation, survival, clustering, phagocytosis, and Aβ plaque compaction^12–15^.

The function of the family member TREM1 (Triggering Receptor Expressed on Myeloid cells-1) is distinct from that of TREM2. In sharp contrast, TREM1 amplifies pro-inflammatory innate immune responses to sterile damage associated molecular patterns (DAMPs) and infectious pathogenic molecular patterns (PAMPs)^16–20^ (**Fig. 1A**). TREM1 synergizes with pattern recognition receptors such as toll-like receptors (TLRs) and NOD-like receptors (NLRs) to increase production of pro-inflammatory cytokines, proteases, and reactive oxygen species (ROS)^21–25^. Although TREM1 has not been identified in GWAS, targeted analysis of the *TREM1* locus has revealed an intronic variant of *TREM1* that is associated with increased amyloid load and rate of cognitive decline^26,27^. Given the pro-inflammatory function of TREM1 and its predominant expression on myeloid cells, we hypothesized that TREM1 may be involved in promoting age-associated, maladaptive innate immune responses relevant to development of AD.

**Figure 1.**
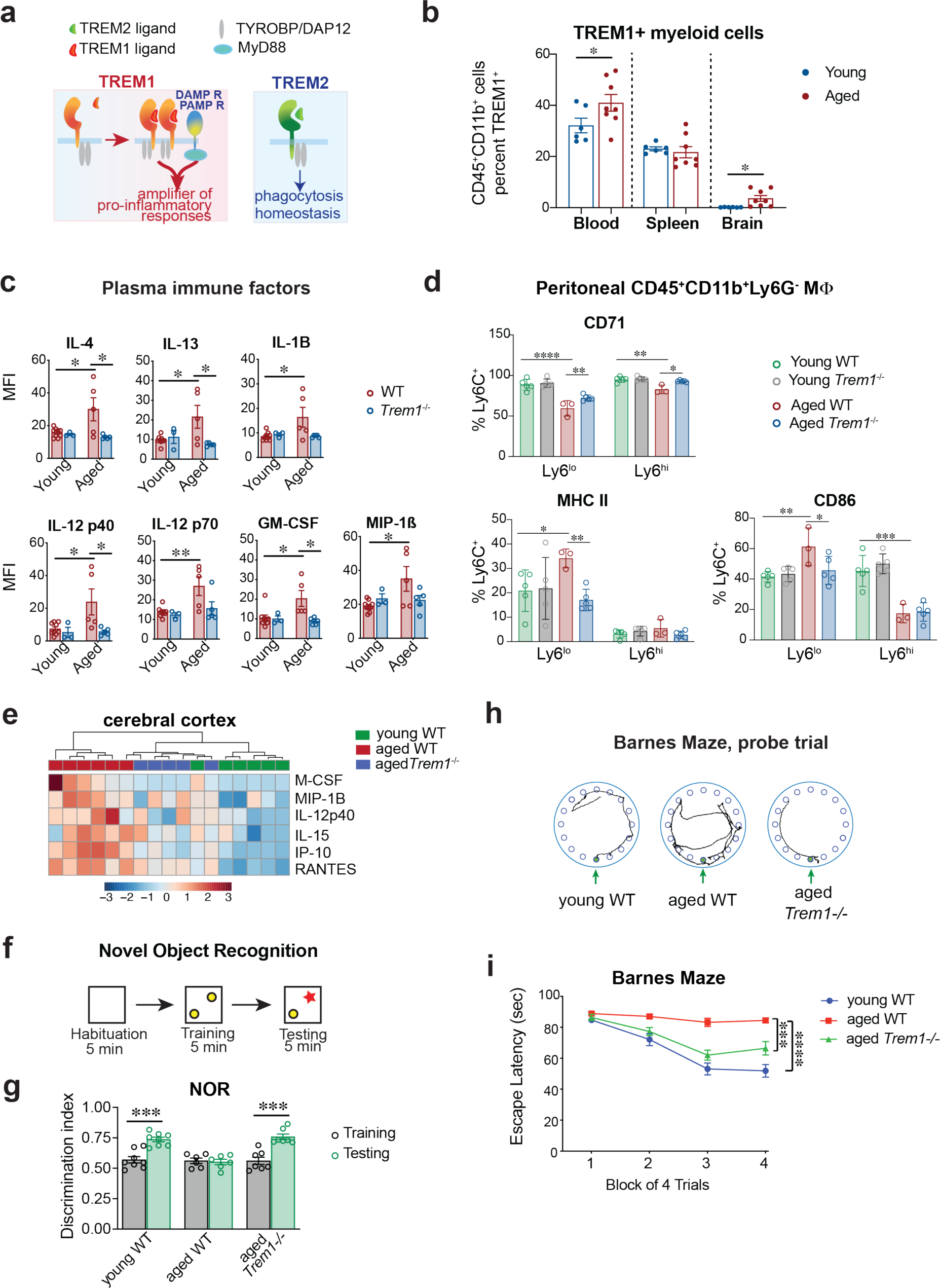
TREM1 deficiency prevents age-associated inflammation and memory decline. a. TREM1 expression on myeloid cells is low under basal conditions, however with stimulation by PAMPs or DAMPs, TREM1 synergizes with innate immune receptors to amplify the pro-inflammatory response. Its functional counterpart TREM2 is anti-inflammatory and promotes phagocytosis and cell survival. b. Percent TREM1 surface expression across blood, spleen and brain CD45^+^CD11b^+^ myeloid cells in young (3 mo) and aged (13-17 mo) mice. Student’s t-test, one-tailed, * *P* < 0.05 (n=6-8 male and female mice per group). c. Multi-analyte quantification of plasma immune factors from young (3 mo) and aged (18.5 mo) WT and *Trem1-/-* mice. Two-way ANOVA with Tukey’s post hoc test, * *P* < 0.05, ** *P* < 0.01 (n=3-8 male mice per group). d. Flow cytometric analysis of anti-inflammatory CD71 and pro-inflammatory MHCII and CD86 in Ly6C^Hi^ and Ly6C^Lo^ peritoneal M<λ, where Ly6C^lo^ and Ly6C^hi^ M<λ correspond to pro- and anti-inflammatory M<λ, respectively, from young 2 mo WT and and aged 22-23 mo WT and *Trem1^-/-^* mice. 2-way ANOVA with post-hoc Tukey multiple comparisons. * *P* < 0.05, ** *P* < 0.01, *** *P* < 0.001, **** *P* < 0.0001 (n=3-5 male mice per group). e. Unsupervised hierarchical clustering of significantly regulated immune factors by one-way ANOVA in cortical lysates from young wild-type (2 mo) and aged WT (25 mo) and *Trem1^-/-^* (23.5 mo) mice. Cluster analysis segregates aged *Trem1^-/-^*mice with young WT mice (n=5-6 male mice per group). f. Procedure for the Novel Object Recognition (NOR) task. Mice were first habituated to testing arena. In the training phase, mice were permitted to freely explore two identical objects. In the testing phase, one object was replaced with a novel object. g. Discrimination index for the NOR task for young WT (5 mo), aged WT (18 mo) and aged *Trem1-/-* (18 mo) mice; paired two tailed t-test (Training vs. Testing), \*\*\**P* < 0.001 (n = 6-8 male and female mice per group). h. Representative movement tracings for the final training trial of the Barnes maze showing rescue effect of *Trem1* deletion. Escape hole is labelled with green arrow. i. Escape latency on the Barnes maze. 2-way repeated measures (RM) ANOVA shows a significant interaction between training and mouse genotype (two-way RM ANOVA interaction *F*(_6,120_) = 7.38, *P* < 0.0001); Tukey’s post hoc test \*\*\**P* < 0.001, \*\*\*\**P* < 0.0001 (n=13-16 male and female mice per group).

Recent studies indicate that myeloid cells, including peripheral monocytes, macrophages (MΦ) and brain microglia, undergo significant bioenergetic decline with aging^28–30^ and this results in loss of beneficial immune responses, including the capacity to resolve inflammation, clear pathogenic substances, and promote a homeostatic anti-inflammatory milieu. Here we report that TREM1 signaling disrupts myeloid cell metabolism and homeostatic immune responses in the contexts of both aging and amyloid accumulation, the two major risk factors for development of AD. We find that TREM1 deficiency restores youthful cellular metabolism and immune function to aging myeloid cells and prevents development of age-associated maladaptive inflammation and hippocampal memory decline in models of aging and AD.

### TREM1 promotes age-associated cognitive decline via the peripheral myeloid compartment

Given that aging is a primary risk factor for development of AD, we first investigated whether TREM1 might affect cognitive aging. TREM1 was robustly expressed in peripheral CD45^+^CD11b^+^ myeloid cells in both blood and spleen and at very low levels in brain microglia (**Supplementary Fig. 1; Fig. 1b**). With aging, TREM1 levels increased in blood (mean 40.96%) and brain (mean 3.63%;). Ex vivo studies of wild type (WT) and *Trem1-/-* peritoneal macrophages (MΦ) confirmed that loss of TREM1 did not alter basal production of immune factors or phagocytosis but significantly dampened the immune response to lipopolysaccharide (LPS) stimulation (**Supplementary Fig. 2a-b**). In vivo, age-associated changes in plasma immune factors were restored to youthful levels in aged *Trem1^-/-^* mice (**Fig. 1c**), as were anti-inflammatory CD71 and pro-inflammatory MHCII and CD86 surface markers in aged Ly6C^lo^ MΦ (**Fig. 1d; Supplementary Fig. 3a-c**). In brain, age-associated increases in chemokines and cytokines also reverted back to youthful levels in aged *Trem1^-/-^*cerebral cortex (**Fig. 1e; Supplementary Fig. 3d**). This rejuvenated immune state in *Trem1-*deficient aged mice was associated with improved memory in the novel object recognition (NOR) task and the Barnes Maze hippocampal spatial memory task (**Fig. f-i; Supplementary Fig. 3e**). These findings indicate that TREM1 promotes development of age-associated inflammatory responses and cognitive decline in aging mice.

To understand whether the observed cognitive rescue in aged *Trem1^-/-^*mice was due to altered brain microglia, we examined transcriptional changes in CD45^lo^CD11b^+^ microglia from young (3 mo) WT and aged (18-20 mo) WT and *Trem1^-/-^* male mice (**Fig. 2a-b; Supplementary Fig. 4a-b**). While there were 9.82% differentially expressed genes (1598 DEGs; FDR, *P*<0.05) in young vs aged WT microglia, there were only 0.08% DEGs (13 DEGs) in aged WT vs aged *Trem1^-/-^* microglia. In sharp contrast, comparison of young WT, aged WT, and aged *Trem1^-/-^* peritoneal MΦ revealed striking differences with 12.9% DEGs in aged WT versus young WT macrophages and 6.54% DEGs in aged *Trem1^-/-^* vs aged WT, but only 2.44% DEGs in the aged *Trem1^-/-^* vs young WT comparisons (**Supplementary Fig. 4c**). The relative paucity of DEGs in the aged *Trem1^-/-^* vs young WT comparison suggests that *Trem1* deficiency restores gene expression to youthful levels in aged MΦ. Indeed, gene expression profiles were largely reciprocally regulated in aged WT compared to both aged *Trem1^-/-^ and* young WT MΦ (**Fig. 2c; Supplementary Fig. 4d-e**). Examination of the chemokine/cytokine pathway (KEGG pathway: mmu04062) revealed that young WT and aged *Trem1^-/-^* transcripts segregated together in reciprocal relation to aged WT macrophages (**Fig. 2d**). In concert with the restoration of youthful pro- and anti-inflammatory surface markers in aged *Trem1^-/-^* peritoneal MΦ (**Fig. 1d**), these findings suggest that the detrimental effects of TREM1 on cognitive aging are mediated predominantly by peripheral myeloid cells and not microglia.

**Figure 2.**
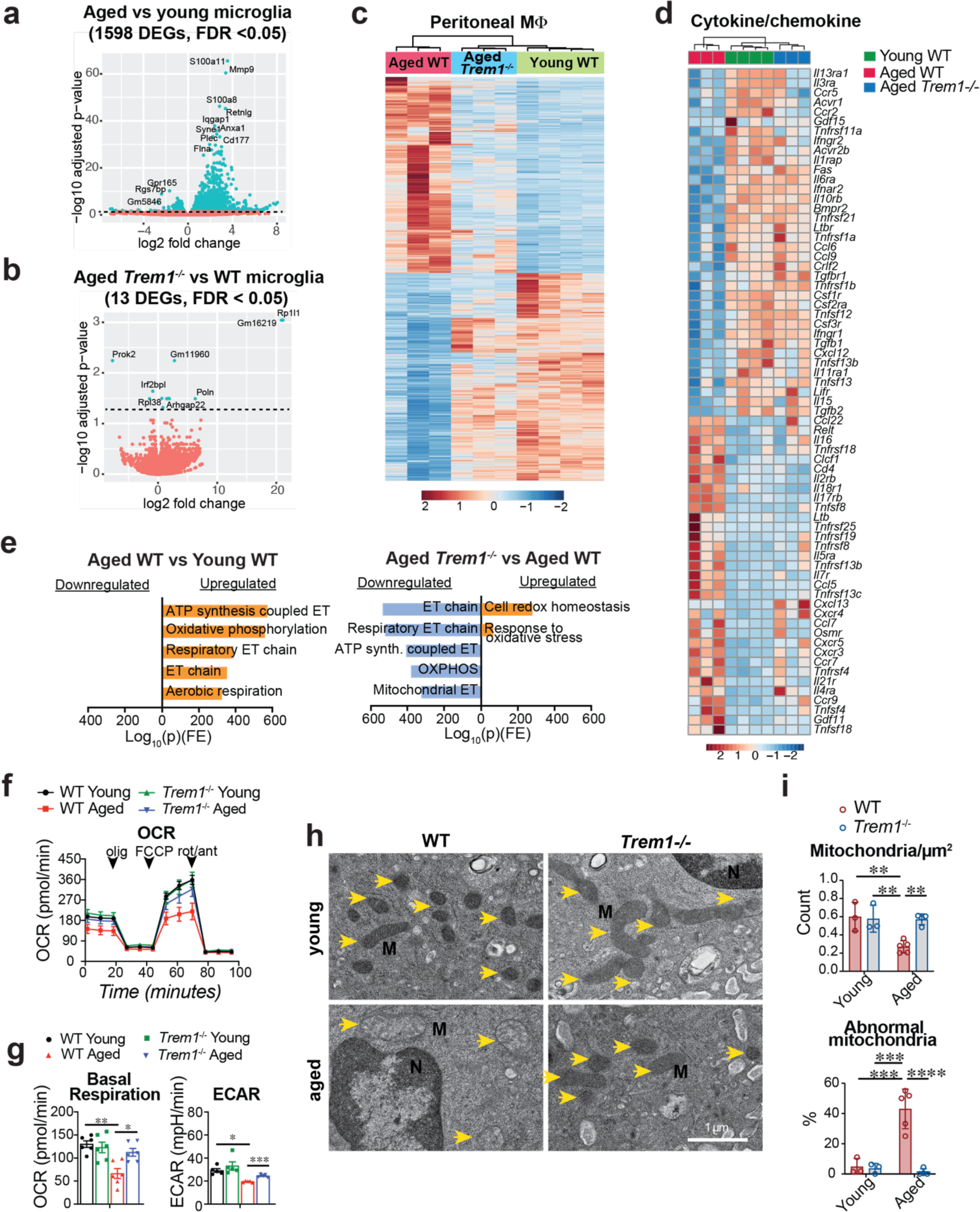
TREM1 is activated in peripheral macrophages in aging. a. Volcano plot of DEGs from aged (18 - 20 mo) vs young (3 mo) CD45^low^CD11b^+^ microglia with -log_10_(*P*) > 1 after FDR adjustment and log_2_ fold change > 1. n=3 flow sorted microglial samples (one sample is pooled from 2 male mice) per age. b. Volcano plot of DEGs from aged *Trem1-/-* vs aged WT microglia with -log_10_(*P*) > 1.25 after FDR and log_2_ fold change > 1. n=3 microglial samples (each sample pooled from 2 male mice) per genotype. c. Unsupervised hierarchical clustering of DEGs from mouse peritoneal M<λ from young WT (2 mo), aged WT (25 mo) and aged *Trem1-/-* (25 mo) mice. Aged *Trem1-/-* macrophage transcriptomic signatures resemble those of young WT macrophages. Scale represents z-score values of FPKM (n=3-4 male mice per group). d. Heatmap (unsupervised hierarchical clustering) of cytokine and chemokines DEGs (*q* < 0.05). Aged *Trem1^-/-^*mice cluster young WT mice. Scale represents z-score values from FPKM (n=3-4 mice per group). e. Pathway enrichment analysis of DEGs in the Mouse MitoCarta3.0 gene list showing reciprocal pathway regulation with *Trem1* deletion in aged macrophages. Scale represents Log(p) x fold enrichment (FE). f. Representative oxygen consumption rate (OCR) traces with SeaHorse for young (4.5 mo) and aged (22-23 mo) wild-type (WT) and *Trem1^-/-^*peritoneal M<λ . Arrowheads indicate application of electron transport-chain inhibitors. Abbreviations: olig: oligomycin, FCCP: carbonyl cyanide-4 (trifluoromethoxy) phenylhydrazone, rot/ant: rotenone/antimycin A. (n=5 male mice per group). g. Quantification of basal respiration and glycolysis (extracellular acidification rate, or ECAR) for young (4.5 mo) and aged (22-23 mo) wild-type (WT) and *Trem1^-/-^* peritoneal M<λ. Two-way ANOVA, Tukey’s multiple comparisons, * *P* < 0.05, ** *P* < 0.01 (OCR, n=6 male mice per group; ECAR, n=5 male mice per group). h. Transmission electron microscopy (TEM) images of young (4.5 mo) and aged (24-26 mo) WT and *Trem1^-/-^* peritoneal M<λ. Yellow arrows indicate mitochondria. Abbreviations: M: mitochondrion; N: nucleus. Scale = 1 µm. i. Numbers of M<λ mitochondria and percent abnormal mitochondria from (**h**). Two-way ANOVA with Tukey’s post hoc test, ** *P* < 0.01 (n= 3-5 male mice per group).

### TREM1 deficiency restores myeloid bioenergetics and nucleotide metabolism

Recent studies indicate that M<λ energy metabolism declines with age, leading to immune activation and pro-inflammatory polarization^28,29^. Accordingly, we performed a targeted analysis of M<λ mitochondria-related genes using Mouse MitoCarta 3.0, a curated mitochondrial gene inventory^31^. This analysis revealed significant age-associated metabolic changes in pathways related to ATP synthesis, electron transport, and oxidative phosphorylation where genes and pathways were restored to youthful levels in aged *Trem1^-/-^* peritoneal M<λ (**Fig. 2e**). Profiling of M<λ bioenergetics demonstrated that deficits in glycolysis (extracellular acidification rate, or ECAR) and mitochondrial oxygen consumption (OCR) were restored to youthful levels in aged *Trem1^-/-^* M<λ (**Fig. 2f-g**). Rejuvenation of mitochondria was confirmed using transmission electron microscopy (TEM) where mitochondrial numbers and integrity were restored in aged *Trem1^-/-^* M<λ to youthful levels (**Fig. 2h-i**). Taken together, transcriptomic, bioenergetic, and ultrastructural data indicate that TREM1 deficiency prevents aging of peripheral M<λ.

To further investigate how TREM1 promotes M<λ bioenergetic decline in aging, we measured 272 metabolic intermediates that are representative of a broad array of cellular metabolic pathways (**Supplementary Table 1**) in young (2 mo) and aged (25 mo) WT and *Trem1^-/-^* peritoneal M<λ. Principal component analysis (PCA) of young WT versus young *Trem1^-/-^* M<λ did not reveal differences between young genotypes (**Supplementary Fig. 5a**). A second study examining metabolic changes between young WT, aged WT, and aged *Trem1^-/-^* peritoneal M<λ showed significant differences between young and aged WT M<λ, and between aged WT and aged *Trem1^-/-^*M<λ, but not between young WT and aged *Trem1^-/-^* M<λ (**Fig. 3a; Supplementary Fig. 5b-c**). *Trem1* deletion restored youthful levels of multiple metabolites, however pathway analysis revealed a prominent rescue of purine, pyrimidine, and NAD^+^ metabolism in aged *Trem1^-/-^* M<λ (**Supplementary Fig. 5d-e**). The common precursor to these three classes of metabolites is ribose-5P, a metabolite generated by the pentose phosphate pathway (PPP) (**Fig. 3b**). Aged *Trem1^-/-^* M<λ showed levels of ribose-5P that were similar to those in young M<λ.

**Figure 3.**
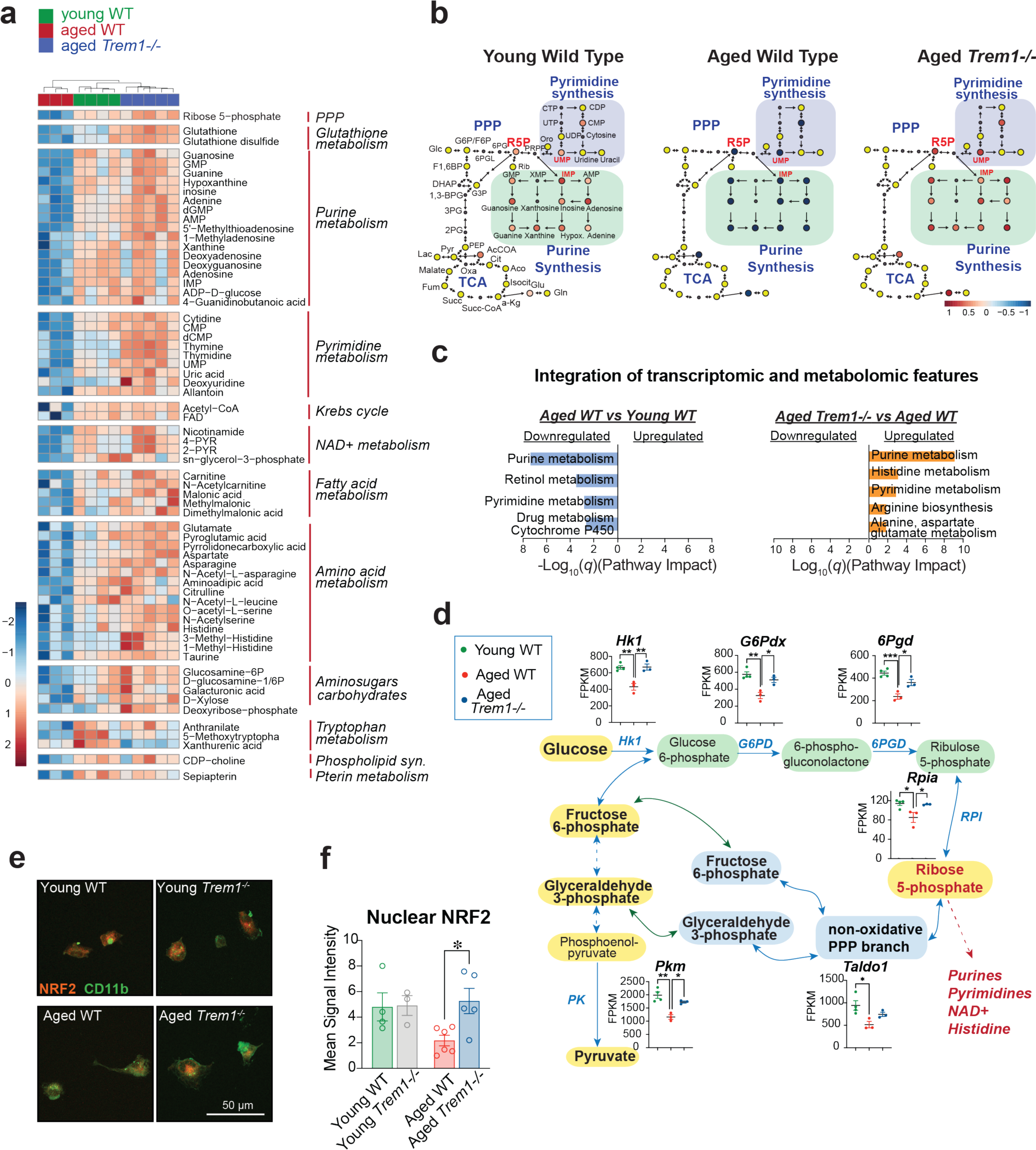
TREM1 suppresses pentose phosphate pathway (PPP) generation of Ribose 5P and purine/pyrimidine synthesis in aged macrophages. a. Unsupervised hierarchical clustering of significantly regulated metabolites from peritoneal macrophages isolated from young WT (2 mo), aged WT (25 mo) and aged *Trem1^-/-^* (25 mo) male mice. TREM1-deficient M<λ cluster with young WT M<λ. Data were analyzed and clustered using MetaboAnalyst 5.0; n=3-5 male mice per group. b. Metabolic pathway map of glycolysis, pentose phosphate pathway (PPP), TCA cycle, and purine and pyrimidine synthesis demonstrates decrease of Ribose 5-Phosphate (R5P), the precursor of purine and pyrimidine synthesis, in aged WT M<λ that is restored to young WT levels with *Trem1* deletion. Colored circles represent z-scored fold change levels for significantly modified metabolites. Yellow circles indicate no significant difference. Small gray circles indicate metabolites that were not measured. c. Integration of transcriptomic and metabolomic features demonstrates enrichment in purine and pyrimidine metabolism. Included in the joint pathway analysis were differentially regulated metabolites (*q* < 0.05) and the highest regulated DEGs (*q-*values < 0.05 and Log2FC > +/-2) and fold change for each feature. No pathways were enriched in the comparison between aged *Trem1^-/-^* and young WT. Data were analyzed using the Joint Pathway Analysis function in MetaboAnalyst 5.0. d. Glycolysis and PPP enzymes (blue) with FPKM quantification. ANOVA followed by Tukey’s post hoc test, \**P* < 0.05, \*\**P* < 0.01, \*\*\**P* < 0.001 (n=3-4 male mice per group). Abbreviations: 6PGD: 6-phosphogluconate dehydrogenase; G6PD: glucose-6-phosphate dehydrogenase; HK1: hexokinase-1, PK: pyruvate kinase; RPI: ribose-5-phosphate isomerase. Enzymes *G6pdx*, *6Pgd*, *Rpia*, *Taldo1* and *Pkm* are known NRF2 target genes. e. Immunofluorescent localization of NRF2 in CD11b^+^ peritoneal M<λ isolated from (4.5 mo) and aged (24-26 mo) WT and *Trem1^-/-^*male mice. f. Quantification of nuclear NRF2 in M<λ from young and aged mice from (**e**). NRF2 signal intensity was calculated within DAPI signal masks for optical slices of 0.5 µm and was averaged across z stacks. 3-5 confocal images were processed per cell using ImageJ. Data analyzed by 2-way ANOVA followed by Tukey’s post hoc test. \**P* < 0.05, (n=3-6 male mice per group).

Integration of transcriptomic data with metabolomic data confirmed a significant effect of TREM1 on purine and pyrimidine synthesis (**Fig. 3c**), which is dependent on sufficient supply of the precursor ribose-5P (**Fig. 3b**). Ribose-5P also serves as a source of glycolytic intermediates through the non-oxidative branch of the PPP, where interconversion of pentose intermediates by transketolases and transaldolases to glycolytic intermediates contributes to pyruvate generation that feeds into the Krebs cycle and mitochondrial respiration. Transcripts encoding enzymes in the oxidative and non-oxidative branches of the PPP, including *G6Pdx*, *6PGd*, *RPI*, and *Taldo1* were reduced in aged WT but restored to youthful levels in aged *Trem1^-/-^* macrophages (**Fig. 3d**). Transcription factor enrichment analysis revealed that the top transcription factor associated with these metabolic changes was NRF2 (**Supplementary Fig. 5f**), which reprograms metabolic transcription to promote nucleotide metabolism^32^ and declines with aging^33^. Indeed, confocal microscopy quantification of nuclear NRF2 in young vs aged peritoneal M<λ demonstrated lower levels of nuclear NRF2 in aged M<λ that were restored to youthful levels in aged *Trem1^-/-^*mice (**Fig. 3e-f; Supplementary Fig. 5g)**. Taken together, metabolomic and transcriptomic data support a role for TREM1 in regulating generation of the critical intermediate ribose-5P through activity of the oxidative and non-oxidative branches of the PPP. Declines in NRF2-dependent gene expression with aging lead to deficits in ribose-5P and replenishment of glycolysis, pyruvate production for mitochondrial respiration, as well as purine/pyrimidine biosynthesis. *Trem1* ablation corrects this deficiency by restoring expression levels of genes encoding PPP enzymes and the glycolytic transcripts hexokinase and pyruvate kinase.

### *Trem1* haploinsufficiency in 5XFAD mice prevents spatial memory deficits

While aging is the primary risk factor for AD, deposition of amyloid is necessary, but not sufficient for progression to AD^34^. We therefore tested whether microglial TREM1 could mediate detrimental effects upon stimulation with oligomeric amyloid ß_42_ (Aß_42_), a highly immunogenic form of amyloid that accumulates in AD (**Fig. 4a-b**). Indeed, TREM1 activation in WT microglia in response to Aß_42_ suppressed mitochondrial oxidative phosphorylation and increased glycolysis, however *Trem1*^-/-^ microglia were completely refractory to the effects of Aß_42_ and maintained homeostatic bioenergetics, reminiscent of the protective effect of *Trem1* deficiency in aging macrophages. We therefore hypothesized that microglial TREM1 may play a role in mediating the neurotoxic effects of locally accumulating amyloid.

**Figure 4.**
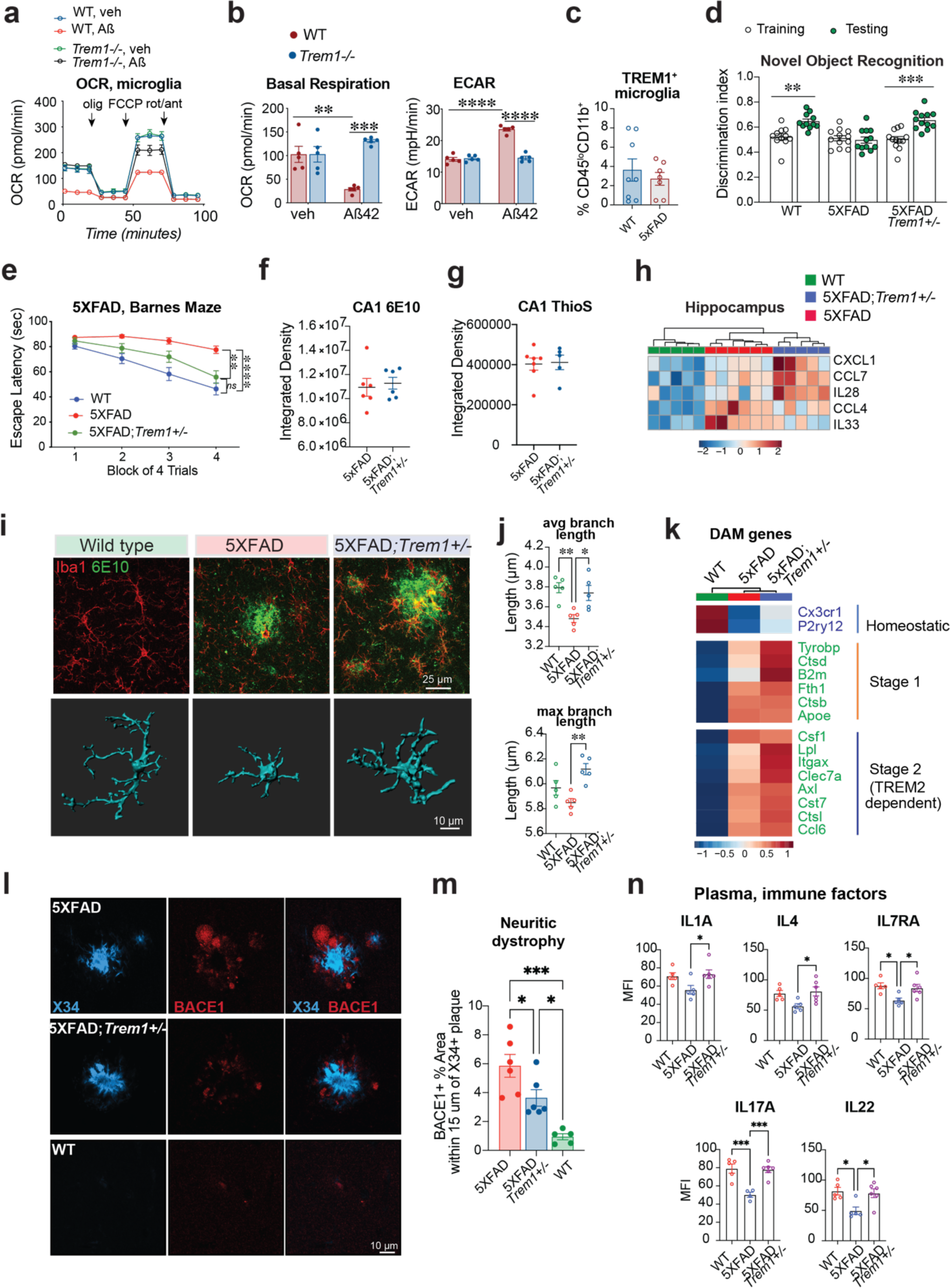
TREM1 deficiency preserves hippocampal function in 5XFAD mice. **a.** Representative oxygen consumption rate (OCR) traces from primary postnatal mouse microglia from WT and *Trem1^-/-^* mice stimulated with vehicle or Aß_42_ oligomers (100 nM) for 20 hours (n=5 wells per genotype/condition). Arrowheads indicate application of electron transport-chain inhibitors. Abbreviations: olig: oligomycin, FCCP: carbonyl cyanide-4 (trifluoromethoxy) phenylhydrazone, rot/ant: rotenone/antimycin A. **b.** Quantification of basal respiration (OCR or oxygen consumption rate) and glycolysis (ECAR or extracellular acidification rate) from **(a)**. Two-way ANOVA with Tukey’s post hoc test, ** *P* < 0.01, *** *P* < 0.001, **** *P* < 0.0001 (n=5 wells per genotype/condition). **c.** Percent TREM1+ microglia in WT vs 5XFAD mice (n= 6-8 male and female mice, 13-17 mo) **d.** Novel Object Recognition (NOR) task was performed on 6-7 mo WT, 5XFAD and 5XFAD;*Trem1^+/-^* mice. Training (clear circles) and testing (green circles) discrimination index differences were assessed using paired *t*-test. Discrimination index of 0.5 indicates chance preference. ** *P* < 0.01, *** *P* < 0.001 (n=11-12 male and female mice per condition). **e.** Escape latency of 5XFAD cohorts on the Barnes maze. Successive training days consisted of a block of 4 trials. Daily mean escape latency was calculated for each mouse. Data were analyzed using 2-way repeated measures (RM) ANOVA with Tukey’s post hoc test. There was a significant interaction between training and mouse genotype (two-way RM ANOVA Interaction *F*(_6,90_) = 4.14, *P* < 0.001). ** *P* < 0.01, **** *P* < 0.0001 (n=10-12 male and female mice per condition). **f.** Mean integrated density of 6E10 signal from CA1 region of hippocampus of 9-10 mo 5XFAD and 5xFAD;*Trem1^+/-^*mice (n=5-7 female mice per group). **g.** Mean integrated density of ThioS signal, a marker of fibrillar amyloid, from CA1 region of hippocampus of 9-10 mo 5XFAD and 5xFAD;*Trem1^+/-^* mice (n=6 female mice per group) **h.** Unsupervised hierarchical clustering of significantly regulated immune factors (by one-way ANOVA) in hippocampal lysates from 9-10 mo WT, 5XFAD, and 5XFAD;*Trem1^+/-^* mice (n=5-6 female mice per group). **i.** Immunofluorescent images of IBA1+ microglia and 6E10 staining of amyloid in the CA1 region of the hippocampus of 9-10 mo WT, 5XFAD, and 5XFAD;*Trem1^+/-^* female mice and 3-dimensional rendering of representative microglia. Scale bar is 25µm (top row) and 10 µm (bottom row). **j.** Quantification of average microglial branch length and maximum branch length from (**i**). One-way ANOVA with Tukey’s post hoc test. * *P* < 0.05, ** *P* < 0.01, **** *P* < 0.0001 (n=5 male mice per condition). **k.** Heatmap depicting group means of significantly regulated DAM signature genes, including homeostatic genes *Cx3cr1*, *P2ry12*, and *Tmem119*, and Stage 1 and Stage 2 genes. Full DAM signature gene set is presented in **Supplementary Fig. 6e**. Individual microglial samples were pooled from two 6-7 mo male mice (n=3 samples per genotype). Scale represents z-score values from FPKM. **l.** Neuritic dystrophy was visualized with anti-BACE1 immunostaining at X34 positive plaques in WT, 5XFAD, and 5XFAD;*Trem1^+/-^* 9-10 mo mice; scale bar=10 µm **m.** Quantification of BACE1 percent area around amyloid plaques, from (**l**). One-way ANOVA with Tukey’s post hoc test, * *P* < 0.05, *** *P* < 0.001 (n= 5-6 female mice per group). **n.** Quantification of plasma immune factors in WT, 5XFAD, and 5XFAD;*Trem1^+/-^* 9-10 mo mice; One-way ANOVA with Tukey’s post hoc test, * *P* < 0.05, *** *P* < 0.001 (n= 5-6 female mice per group).

Flow cytometry of microglia in the 5XFAD model^35^ of amyloid accumulation showed low TREM1 surface expression that did not differ from age-matched WT controls (**Fig. 4c**). Similarly, in the *APP^Swe^PS1^ΔE^*^9^ model^36^ of amyloid accumulation, immunostaining of TREM1+ Iba1+ microglia did not show a significant induction of TREM1 expression compared to WT (**Supplementary Fig. 6a**), suggesting that microglial TREM1 expression is not significantly changed in the context of accumulating amyloid. To determine whether TREM1 might nevertheless promote cognitive deficits in the context of amyloid and independently of age, we examined 6-7 mo 5XFAD mice that were heterozygous for *Trem1*. *Trem1* haploinsufficiency in 5XFAD mice prevented cognitive deficits in both the novel object recognition (NOR) memory test and Barnes maze test of spatial learning and memory (**Fig. 4d-e**). While no differences in levels of dense core or diffuse amyloid plaque were detected (**Fig. 4f-g; Supplementary Fig. 6b**), several immune factors were changed between 5XFAD and 5XFAD;*Trem1^+/-^*hippocampi (**Fig. 4h; Supplementary Fig. 6c**). Examination of microglial morphology revealed a striking preservation of resting characteristics in 5XFAD;*Trem1^+/-^* mice, where microglia remained homeostatic and not activated, with average and maximal branch lengths indicative of a surveilling phenotype (**Fig. 4i-j; Supplementary Fig. 6d**).

To investigate the molecular mechanisms underlying this preservation of homeostatic morphology, we examined transcriptional differences in CD45^lo^CD11b^+^ microglia isolated from 6-7 mo male WT, 5XFAD and 5XFAD;*Trem1^+/-^* mice. We observed the expected transition from WT to the disease-associated microglial signature^15^ (DAM) in 5XFAD microglia, but there were no significant differences between 5XFAD and 5XFAD; *Trem1^+/-^* DAM signatures (**Supplementary Fig. 7a-c**). Although not significant, there was a trend in 5XFAD;*Trem1+/-* microglia towards higher homeostatic transcript levels and selected Stage 1 and Stage 2 phagocytic/inflammatory DAM genes^15^ (**Fig. 4k**). Consistent with the muted effect on the DAM signature, we did not observe significant changes by flow cytometry in homeostatic markers CX3CR1 and Tmem119 between 5XFAD and 5XFAD;*Trem1+/-* microglia (**Supplementary Fig. 7d-e**). However, quantification of BACE1 immunostaining at amyloid plaques, a marker of neuritic dystrophy^37^, was significantly reduced in 5XFAD;*Trem1^+/-^* as compared to 5XFAD mice, suggesting a neuroprotective effect of *Trem1* haploinsufficiency (**Fig. 4l-m**). We also identified a restoration to WT levels of several plasma cytokines in 9-10 mo 5XFAD;*Trem1^+/-^* mice (**Fig. 4n**), suggesting a peripheral effect of *Trem1* haploinsufficiency. Taken together, these data suggest a protective role of TREM1 deficiency that is independent of amyloid and microglial transcriptional responses to amyloid.

### TREM1 deficiency in aged *APP^Swe^* mice rescues spatial memory deficits and cerebral glucose metabolism

We then investigated the functional consequences of TREM1 activity in a second model, the TG2576 *APP^Swe^* model of amyloid accumulation^38^. *APP^Swe^*mice develop progressive hippocampal memory deficits beginning at 6 months of age and amyloid plaques after 12 months of age. We examined the role of TREM1 in the interaction of aging and amyloid deposition, the context in which AD develops. In the novel objection recognition test, memory was disrupted in aged 18-21 mo male and female *APP^Swe^* mice compared to WT littermates, however loss of one or both *Trem1* alleles restored memory to WT levels (**Fig. 5a**). In the Barnes maze, spatial memory was significantly rescued in *APP^Swe^;Trem1^-/-^* mice as compared to WT littermates (**Fig. 5b**; **Supplementary Fig. 7e**). As in the 5XFAD model, this behavioral rescue occurred in the absence of changes in amyloid deposition (**Fig. 5c-d**). Selected hippocampal immune factors were increased in *APP^Swe^* with variable effects of *Trem1* deficiency; however in plasma, increases in inflammasome products IL1α and IL1ß were elevated in *APP^Swe^*mice but restored to WT levels with *Trem1* deficiency (**Fig. 5e-f**). These data indicate that TREM1 promotes a systemic pro-inflammatory state in the context of aging and accumulating amyloid.

**Figure 5.**
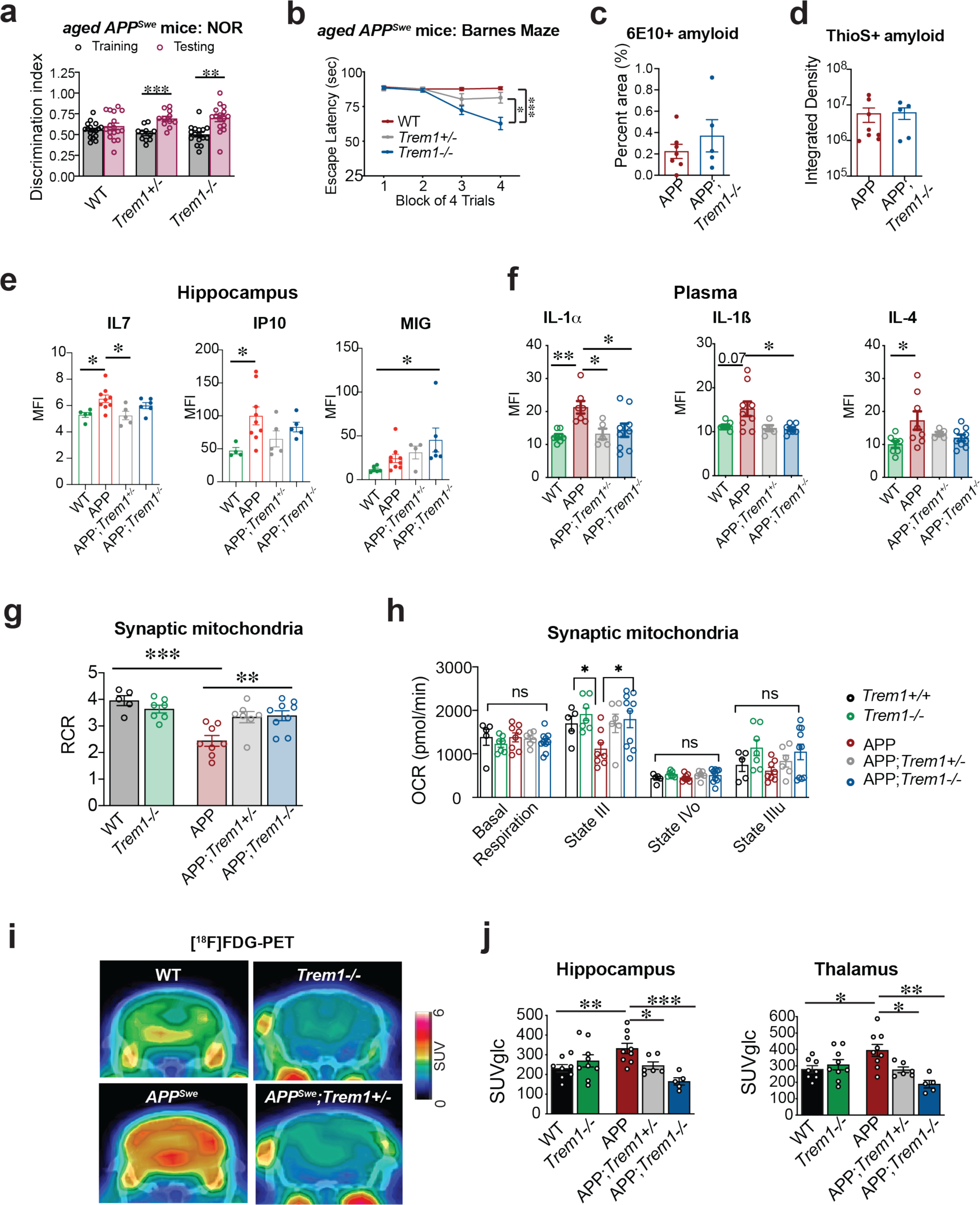
TREM1 deficiency preserves hippocampal spatial memory and brain glucose metabolism in *APP^Swe^* mice. **a.** Discrimination index during the NOR task for aged (18-21 mo) *APP^Swe^*, *APP^Swe^/Trem1^+/-^*and *APP^Swe^/Trem1^-/-^* mice; paired t-test of Training vs. Testing. ** *P* < 0.01, *** *P* < 0.001 (n=11-17 male and female mice per group). **b.** Escape latency of *APP^Swe^*cohorts in the Barnes maze. Daily mean escape latency was calculated for each mouse and mouse groups were analyzed using 2-way repeated measures (RM) ANOVA with Tukey’s post hoc test. There was a significant interaction between training and mouse genotype (two-way RM ANOVA Interaction *F*(_6,132_) = 12.24, *P* < 0.0001). * *P* < 0.05, *** *P* < 0.001 (n=14-17 male and female mice per condition). **c.** Percent area positive for 6E10 immunostaining in hippocampus of 18-21 mo *APP^Swe^* and *APP^Swe^*;*Trem1^-/-^* mice. Student’s two tailed t-test (n=5-8 male/female mice per group). **d.** Average integrated density for ThioS^+^ signal in hippocampus of 18-21 mo APP*^Swe^* and *APP^Swe^*;*Trem1^-/-^* mice. Student’s two tailed t-test, n=5-8 male/female mice per group. **e.** Significantly regulated hippocampal immune factors in 20-23 mo in WT, *APP^Swe^*, *APP^Swe^/Trem1^+/-^* and *APP^Swe^/Trem1^-/-^* mice. One-way ANOVA with Tukey’s post hoc test, * *P* < 0.05 (n=4-9 female mice/group). **f.** Significantly regulated plasma cytokines in 20-23 mo in WT, *APP^Swe^*, *APP^Swe^/Trem1+/-* and *APP^Swe^/Trem1-/-* mice. One-way ANOVA with Tukey’s post hoc test, * *P* < 0.05, ** *P* < 0.01 (n=4-9 female mice/group). **g.** Respiratory control ratio (RCR) measured in isolated synaptic mitochondria fractions from 18-24 mo mouse brain. High RCR indicates mitochondria with a high capacity for substrate oxidation and ATP turnover and a low proton leak. One-way ANOVA with Tukey’s post hoc test. ** *P* < 0.01, *** *P* < 0.001 (n=5-9 male mice per condition). **h.** Synaptic mitochondria OCR (pmol/min) calculated basally and for state III, state IVo, and state IIIu in 18-24 mo in WT, *APP^Swe^* and in *APP^Swe^;Trem1^-/-^* mice. State III reflects maximal ADP-stimulated respiration and State IV reflects the return to a basal state of respiration after addition of ATP synthase inhibitor oligomycin. ANOVA with Tukey’s post-hoc test. \**P* < 0.05 (n=5-9 male mice per condition). **i.** Representative coronal brain [^18^F]FDG-PET/CT images of 17-19 mo mice showing normalization of cerebral glucose metabolism in *APP^Swe^/Trem1+/-* female mice. SUV: standardized uptake values. Static 20 min PET images were acquired at 75-95 min following [^18^F]FDG injection. **j.** Quantification of [^18^F]FDG-PET signal in hippocampus and thalamus. SUVs were normalized to individual blood glucose concentrations. One-way ANOVA with Tukey’s post hoc test; * *P* < 0.01, ** *P* < 0.01, *** *P* < 0.001 (n=5-9 female mice per condition).

Healthy neuronal mitochondrial function is critical for neurotransmission and circuit integrity, so we investigated whether myeloid TREM1 regulates bioenergetic function of synaptic mitochondria isolated from synaptosome fractions of *APP^Swe^* mice lacking one or both *Trem1* alleles. We determined the extent of coupling between the electron transport chain (ETC) and oxidative phosphorylation of ADP to ATP in synaptosomes^28,39^. The respiratory control ratio (RCR), represented as the ratio of State III/State IVo, was significantly reduced in 18-24 mo *APP^Swe^* mice as compared to age-matched WT mice, consistent with poor neuronal mitochondrial function (**Fig. 5g-h**). However, loss of one or both *Trem1* alleles restored synaptic mitochondrial function to WT levels. The improved coupling of electron transport and ATP synthesis suggests that in the setting of accumulating amyloid, *Trem1*-deficient microglia maintain homeostatic immune responses that preserve healthy synaptic mitochondrial function.

Using an orthogonal approach to measure neuronal metabolic function, we assessed cerebral glucose metabolism using positron emission tomography (PET) and the glucose analogue tracer 2-deoxy-2-^18^F fluoro-D-glucose ([^18^F]FDG PET) in 17-19 mo female *APP^Swe^* mice. Although [^18^F]FDG PET reliably demonstrates hypometabolism in human parietal cortex in patients with AD, in transgenic mutant APP models, [^18^F]FDG PET signal can vary depending on experimental factors such as anesthesia, type of APP mutation, and age^40^. Here, compared to wild type controls, *APP^Swe^* hippocampus and thalamus showed a significant increase in glucose uptake that was prevented with deletion of one or both *Trem1* alleles (**Fig. 5i-j).** The normalization of brain glucose metabolism with partial or complete *Trem1* deficiency suggests that accumulating amyloid in *APP^Swe^* mice triggers TREM1-dependent microglial responses that disrupt brain glucose metabolism. Together, these results suggest that *Trem1* deficiency preserves cellular and cerebral bioenergetics in aging *APP^Swe^*mice.

### TREM1 increases in Alzheimer’s Disease

Mendelian randomization (MR) is a method for inferring causal associations between genetic variants and disease. A recent study applying MR to plasma proteins found that increased plasma levels of the soluble ectodomain of TREM1, or sTREM1, was associated with increased AD risk^41^. We validated this finding independently using plasma proteins measured using a multiplexed, aptamer-based approach (SOMAscan assay) in 35,559 Icelanders^42^ to determine which variants associated with plasma sTREM1 and sTREM2. We then assessed AD risk associated with these variants in a second cohort of 75,024 cases and 397,844 controls from the UK Biobank^43^. These MR associative analyses indicated that both sTREM1 and sTREM2 levels were causally associated with AD risk (**Fig. 6a and Supplementary Fig. 8a-b**). Increased sTREM1 levels were associated with increased AD risk and conversely, increased sTREM2 levels were associated with decreased AD risk.

**Figure 6.**
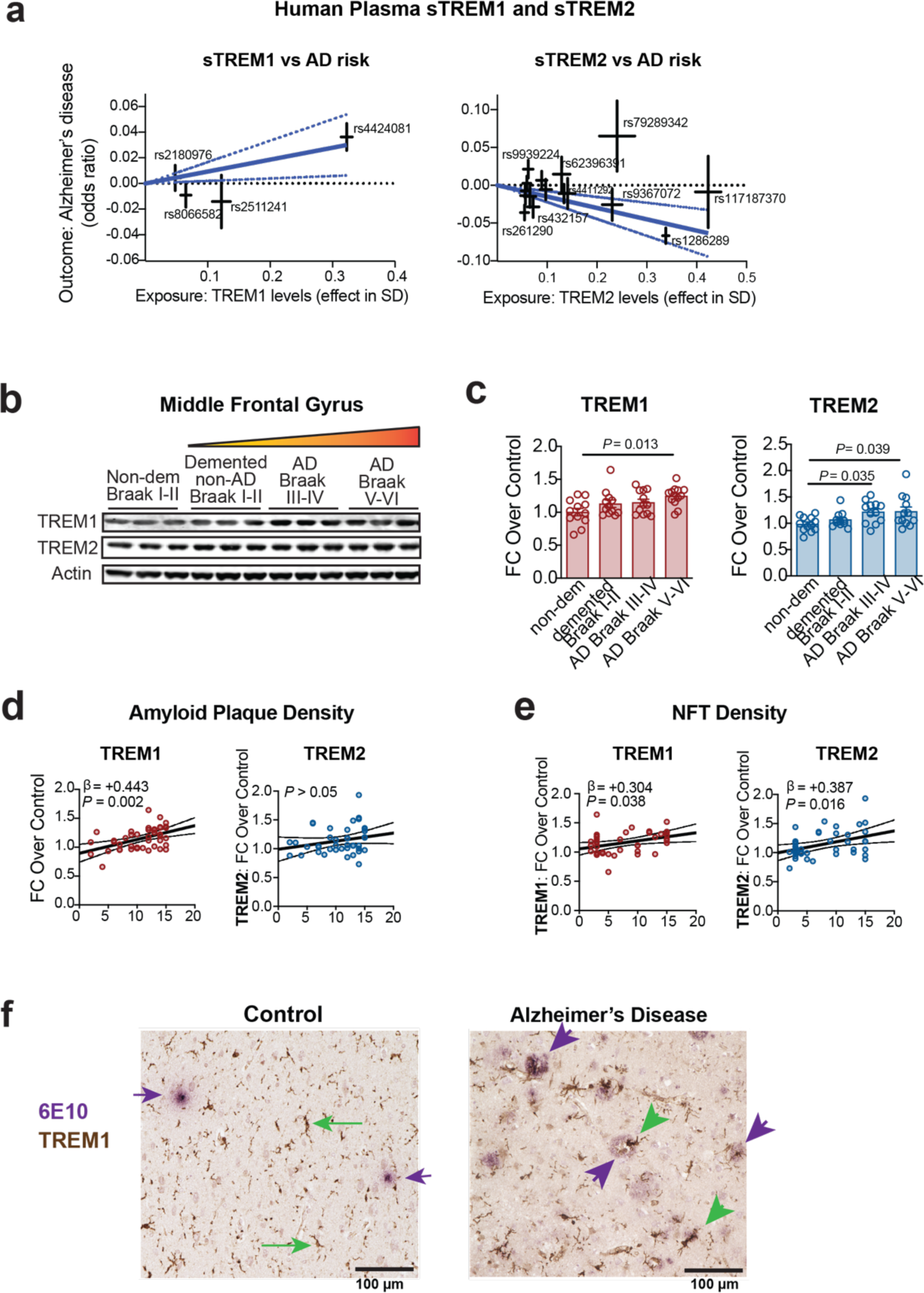
TREM1 is an AD risk gene and expression positively associates with increasing severity of pathology. **a.** Mendelian Randomization (MR) analysis of relationship between plasma levels of soluble TREM1 (sTREM1) and sTREM2 (exposure) and Alzheimer’s disease risk (outcome). Blue lines are estimated MR-inverse variance weighted effects, and dashed lines indicate the 95% confidence interval for MR effects. Increased plasma sTREM1 level was associated with increased AD risk (β = +0.0929, [95% CI = 0.019 to 0.166] *P* = 0.013), while increased plasma sTREM2 level was associated with decreased AD risk (β = -0.15 [95% CI = -0.223 to -0.077] *P* = 5.61×10^-05^). **b.** Representative immunoblot of TREM1 and TREM2 proteins in the middle frontal gyrus across clinicopathological diagnoses: non-demented Braak I-II, demented non-AD Braak stages I-II, AD Braak III-IV and AD Braak V-VI (n=12 donors per group). **c.** Quantification of TREM1 and TREM2 protein expression, normalized to ß-actin and expressed as fold-change over non-demented control. Group differences were analyzed using ANCOVA followed by Tukey’s post hoc test. Age and sex were included as covariates. Each point represents an individual donor (n=12 donors per group). **d.** Linear regression analyses between amyloid plaque and neurofibrillary tangle (NFT) density and TREM1 (left plot) and TREM2 (right plot) levels determined by quantitative immunoblotting. Note lack of significant correlation for TREM2. Model was adjusted for age and sex. Plotted are the 95% confidence bands of the best-fit line from the linear regression. The standardized regression coefficients (ß) and P-values from the linear model are shown. n=43 donors (amyloid plaque density vs TREM1) and n=41 donors (NFT density vs TREM1). **e.** Linear regression analyses between neurofibrillary tangle (NFT) density and TREM1 (left plot) or TREM2 (right splot) levels determined by quantitative immunoblotting. Note significant correlation of both TREM1 and TREM2 with NFT levels. Model was adjusted for age and sex. Plotted are the 95% confidence bands of the best-fit line from the linear regression. The standardized regression coefficients (ß) and P-values from the linear model are shown. n=43 donors (amyloid plaque density vs TREM1) and n=41 donors (NFT density vs TREM1). **f.** Double immunostaining for TREM1 (brown) and 6E10 (purple) show expression of TREM1 in microglia in control and AD superior temporal cortex. Incidental small 6E10^+^ amyloid deposits (horizontal purple arrows) are associated with highly ramified TREM1^+^ microglia (green horizontal arrows) in control brain; in AD brain, amyloid plaques are abundant and compact and are associated with activated TREM1^+^ microglia (purple and green diagonal arrows, respectively). Scale bar = 100 μm.

To probe the relevance of TREM1 to AD pathology, we investigated its expression in human post-mortem brain that harbored increasing levels of Braak pathology, the basis for neuropathological diagnosis of AD^44^. We examined TREM1 levels in middle frontal gyrus, or BA 9/46^45^, a brain region that in AD demonstrates a severe bilateral gray matter loss^46^, synaptic loss, and high Aβ burden^47^. The tissue was obtained from subjects classified as: non-demented (Braak I–II, n = 12), demented Braak I–II (non-AD, n = 12), AD Braak III–IV (AD-mid, n = 12) and AD Braak V–VI (AD-high, n = 12). Diagnostic groups were balanced for age, gender, and post-mortem interval. Demographic information of the cohort is shown in **Supplementary Table 2**. Quantification of protein levels by immunoblotting revealed a steady increase of both TREM1 and TREM2 in frontal cortical lysates from non-demented, demented Braak I-II, AD-Braak III-IV, and AD-Braak V-VI donors (**Fig. 6b-c and Supplementary Fig. 8c-d)**. The fold change increase over control was identical for TREM1 and TREM2 at 1.24-fold. Thus, both TREM1 and TREM2 increase with clinical AD progression, reaching significance at the AD-high stage which is characterized by extensive neocortical involvement of neurofibrillary tangles (NFTs) along with high Aß plaque burden in the frontal cortex. TREM1 levels were positively associated with both amyloid and tau pathology, whereas TREM2 levels were positively associated with tau pathology alone (**Fig. 6d-e**). The relationship between TREM2 and tau is supported by cerebrospinal fluid (CSF) soluble TREM2 (sTREM2) biomarker data, where increased levels of CSF sTREM2 are observed in patients and controls with increased tau pathology^48,49^.

To determine the cellular localization of TREM1, we carried out TREM1 immunostaining in Iba1+ microglia from both control and AD superior temporal cortex. TREM1 expression in AD brain localized to morphologically activated microglia near 6E10 positive amyloid deposits, in contrast to control brain, where TREM1+ microglia exhibited thin and ramified processes (**Fig. 6f)**. Immunofluorescent staining of TREM1 in post-mortem AD frontal cortical free-floating sections revealed co-localization of TREM1 with CD163, a scavenger receptor expressed in perivascular macrophages and activated microglia^50^ (**Supplementary Fig. 8e**). Together, the above findings indicate that TREM1 expression increases with increasing amyloid and tau burden and positively associates with AD risk.

## DISCUSSION

Although significant interest has centered on the protective microglial phenotype promoted by TREM2 in amyloid accumulation in AD, our findings point to its functional counterpart, TREM1, as a potent driver of age- and amyloid-dependent myeloid dysfunction and cognitive decline. In the setting of physiological aging, transcriptomics indicate that peripheral myeloid TREM1, and not microglial TREM1, drives age-associated inflammation and cognitive decline. Metabolomic analysis revealed that with aging, basal TREM1 expression suppressed glucose metabolism, particularly the generation of ribose-5P that is critically required for both glycolysis and biosynthesis of purines, pyrimidines, and NAD+. TREM1-mediated metabolic suppression was driven in part by decreased expression of genes encoding critical enzymes in the oxidative and non-oxidative PPP that are regulated by NRF2. In contrast, preclinical modeling in transgenic models of amyloid accumulation, a necessary event for development of AD^34^, revealed a TREM1-dependent disruption of spatial memory that was associated with both peripheral and microglial immune changes. The phenotypic rescue from TREM1 deficiency was independent of amyloid accumulation, consistent with the finding that TREM1 deficiency rendered microglia refractory to Aß_42_-induced changes in bioenergetics. Although WT microglia responded by increasing aerobic glycolysis and suppressing mitochondrial oxidative phosphorylation, consistent with the Warburg response^51^, the bioenergetics of *Trem1-*deficient microglia remained unperturbed. In 5XFAD mice, *Trem1* haploinsufficiency preserved homeostatic microglial morphology and reduced neuritic dystrophy, a rescue that occurred in the absence of changes in the DAM signature, a transcriptional pattern that reflects microglial responses to accumulating amyloid. In aging *APP^Swe^* mice, loss of one or both *Trem1* alleles also prevented cognitive decline independently of amyloid and elicited a neuroprotective effect by normalizing synaptic mitochondrial integrity and cerebral glucose uptake. In human subjects, we demonstrated that increasing TREM1 expression in postmortem brain associated positively with increasing amyloid and tau pathologies in AD subjects. Thus, TREM1 functions as a negative regulator of myeloid bioenergetics and homeostatic immune responses and promotes cognitive decline in both aging and amyloid accumulation.

Flow cytometry measurements demonstrated relatively low levels of surface TREM1 expression in microglia as compared to peripheral myeloid cells in young and aged mice. This would suggest a minimal contribution of microglial TREM1 to disrupted neuronal function in aging. This conclusion was supported by transcriptomic analyses in *Trem1*-deficient mice, which showed significant changes in peripheral macrophages but negligible changes in aging microglia, and metabolomic analyses of peripheral macrophages, where youthful bioenergetics were maintained in aged *Trem1*-deficient mice. The mechanisms by which *Trem1*-deficient peripheral myeloid cells regulate neuronal function may include beneficial conditioning of the blood and blood-brain barrier from improved macrophage function throughout the periphery.

In 5XFAD mice, there was also no significant induction of TREM1 surface expression on microglia, a finding in line with the muted microglial transcriptional response to *Trem1* haploinsufficiency. However, microglia in 5XFAD;*Trem1^+/-^* mice demonstrated a marked resilience by maintaining their homeostatic morphology; here, *Trem1* haploinsufficiency led to persistence of a non-activated, surveilling phenotype. A potential explanation may reside in the finding that *Trem1*-deficient microglia are refractory bioenergetically to the immunogenic effects of amyloid-ß and therefore will not launch immune responses that are neurotoxic. It is possible that microglial TREM1, even at low expression levels, mediates highly neurotoxic effects, a possibility that is supported by the significant decrease in neuritic dystrophy in 5XFAD;*Trem1^+/-^* mice. However, given the restoration to WT levels of plasma cytokines in 5XFAD;*Trem1^+/-^* mice, we cannot exclude a peripheral neuroprotective effect of *Trem1* haploinsufficiency in addition to a protective microglial effect.

In the *APP^swe^* model of amyloidosis, immune activation is linked to increased brain metabolism as measured by glucose uptake visualized by PET. While it remains unclear what cell type is driving the PET signal in AD mice, our FDG-PET findings are in line with a single-cell radiolabeling approach indicating that microglia can be a major contributor to FDG uptake in AD PS2APP and APPPS1 mouse models of amyloidosis^52^. Data suggest that microglia display higher glucose uptake than neurons and astrocytes, although there may be methodological explanations for these findings^53^. Our analyses of synaptic mitochondrial bioenergetics in *APP^swe^* mice showed prominent deficits – prevented in *Trem1^+/-^ and Trem1^-/-^* genetic backgrounds – that could potentially lead to compensatory increases in glucose uptake in *APP^swe^* mice. However other cell types, particularly astrocytes, may also be driving glucose uptake. Thus, the normalization of the FDG-PET signal with loss of one or both *Trem1* alleles may reflect changes in additional cell types directly and indirectly affected by TREM1-mediated changes in microglia. Overall, our findings support the emerging concept that cellular metabolism – in microglia and peripheral macrophages – is a major regulator of immune responses that regulate cognitive function in aging and neurodegeneration.

Systemic and brain inflammation are major contributing factors to the initiation and progression of age-associated cognitive decline and neurodegeneration. Our findings demonstrate a highly deleterious role of basal TREM1 signaling in aging driven by its disruption of myeloid glucose and nucleotide metabolism. In the aging brain, and in the aging brain that is accumulating amyloid, TREM1-mediated disruption of homeostatic and youthful bioenergetics promotes immune responses that are neurotoxic. These findings suggest an alternative approach to diseases of aging, where myeloid cells might be reprogramed to a healthier metabolic phenotype, providing a critical disease-modifying effect needed to slow or halt progression to AD.

## Acknowledgements

This work was supported by RF1AG053001K (KIA), RO1NS100180 (KIA), RF1AG058047 (KIA), RF1AG070839 (KIA), 1P50 AG047366 (KIA), American Heart Foundation/Allen Frontiers Award (KIA), The Weston Havens Foundation (KIA), The Archer Foundation (KIA), Stanford School of Medicine Dean’s Postdoctoral Fellowship (ENW), HHMI Hanna H. Gray Fellows Program (MRM), Burroughs Wellcome Fund PDEP (MRM), Alzheimer’s Association Research Fellowship (KAZ), The Paul and Daisy Soros Fellowship for New Americans (PSM), the Gerald J. Lieberman Fellowship (PSM), Marie Skłodowska-Curie Grant 888494 (EB), Stanford School of Medicine Dean’s Postdoctoral Fellowship (EB). Data and tissue used in the preparation of this article were obtained from the Arizona Study of Aging and Neurodegenerative Disorders (AZSAND) (www.brainandbodydonationprogram.org), supported by the National Institute of Neurological Disorders and Stroke (U24 NS072026), the National Institute on Aging (P30 AG19610), the Arizona Department of Health Services, the Arizona Biomedical Research Commission and the Michael J. Fox Foundation for Parkinson’s Research. We are grateful to the UW Alzheimer’s Disease Research Center, Pacific Northwest Udall Center, and the Department of Veterans Affairs for providing brain samples, to the Stanford Cyclotron & Radiochemistry Facility staff who produced 18F FDG, especially Drs. Murugesan Subbarayan, Francis Balmaceda Jr, and Ka Chun Ho, and the SCi3 small-animal imaging facility, Stanford Shared FACS Facility, the Stanford Human Immune Monitoring Core, and the Stanford Neuroscience Microscopy Service. KIA is a Chan Zuckerberg-San Francisco Biohub Investigator.

## AUTHOR CONTRIBUTIONS

E.N.W. and K.I.A. conceived and planned this study. E.N.W., T.G.B., M.D.G., M.L.J., M.S.B, J.D.R. and K.I.A contributed to supervising experimental design. E.N.W., C.W., H.E.E., M.S.S., M.R.M., Q.W., K.A.Z., A.C., J.A.R.B., E.G., P.S.M., E.B., Y.J.T., C.A.I., Y.L.G., M.P., H.C., P.J., Q.L., S.S.M., A.J.Z., M.X., J.U., J.H., A.S.D. and G.E.S. conducted the experiments. E.N.W., C.W., H.E., M.S.S., A.C., J.A.R.B., E.G., P.S.M., Y.L.G., A.J.Z. M.X., J.U., and J.H. performed collection of the data and statistical analysis. E.N.W. and K.I.A. wrote the manuscript. All authors reviewed and approved of the manuscript.

## COMPETING FINANCIAL INTRESTS STATEMENT

M.L.J and K.I.A are co-founders and Scientific Advisory Board members of Willow Neuroscience, Inc.

