## Supplementary data for "TREM1 disrupts myeloid bioenergetics and cognitive function in aging and Alzheimer’s disease models"

**Supplementary Materials**

I. **Methods**

II. **Supplementary Figures**

III. **References**

### MATERIALS AND METHODS

#### Animals

This study was conducted in accordance with National Institutes of Health (NIH) guidelines and the Institutional Animal Care and Use Committee at Stanford University approved protocols. *Trem1*$^{-/-}$ mice on a C57BL/6 genetic background have been previously described[10] and were crossed with either 5XFAD[11] (Jackson Labs) or *APP*$^{Swe}$ [12] mice (Taconic) or with C57BL/6J mice (Jackson Labs). All mice were housed in an environment controlled for lighting (12 hour light/dark cycle), temperature, and humidity, with food and water available *ad libitum.* The Stanford Veterinary Service Center monitors and maintains pathogen-free mouse housing as described in http://med.stanford.edu/vsc/about/rodent-handbook/rodent-diseases.html.

#### Flow cytometry of microglia, peripheral myeloid cells, and peritoneal macrophages

Mice were terminally anaesthetized. Blood and splenocytes were collected and lysed with ACK lysis buffer for 10 min at room temperature. All cells were then resuspended in FACS buffer. Dead cells were excluded by staining with Live/dead Aqua (Thermo Fisher). Cells were stained with antibodies for 30 min on ice. Microglial purity was confirmed by CD11B-A700 (Thermo Fisher, clone M1/70), CD45-BUV395 (BD Biosciences, clone 104), Gr1-V450 (Thermo Fisher, clone RB6-8C5), and CD206-BV605 (Biolegend, clone C068C2). Microglial homeostasis was examined by CX3CR1-PE/Cy7 (Biolegend, clone SA011F11), P2RY12-PE (Biolegend, clone S16007D), and Tmem119-FITC (Thermo Fisher, clone V3RT1GOsz). TREM1 expression in microglial was examined by Trem1-APC (R&D, clone 174031). Cells were centrifuged, washed, and resuspended in FACS buffer for flow cytometry on an LSR II (BD Biosciences) with FACSDiva software (BD Biosciences). FlowJo software was used for the analysis and depiction of the gating strategy.

Peritoneal macrophages were collected from WT and *Trem1*$^{-/-}$ mice. The peritoneal space was flushed with 10 mL ice-cold PBS. Collected cells were added to 1 mL of ACK lysis buffer (Cat# A1049201, Thermo Fisher Scientific) for 5 minutes at room temperature. The prepared samples were then washed with FACS buffer (500 mL PBS, 1g BSA, 1mL 5M EDTA) and centrifuged at 500 g for 3 minutes. Cells were resuspended in FACS buffer containing 1:200 of primary antibodies: CD11b-BV421 (Cat# 101235, Biolegend), Ly6G-BV605 (Cat# 563005, Biolegend), LY6C-V450 (Cat# 560594, Biolegend), CD86-APC (Cat# 105012, Biolegend), CD45-AF700 (Cat# 560510, Biolegend), MHCII-FITC (Cat# 11-5321-82, Thermo Fisher Scientific), CD71-PerCP/Cyanine5.5 (Cat# 113816, Biolegend), Live/Dead-Aqua (Cat# L34957, Thermo Fisher Scientific), and incubated for 20 minutes at 4°C. Cells were centrifuged at 500 g for 3 minutes, washed one time with FACS buffer, and resuspended in 4% PFA for 25 minutes at 4°C. After fixation, cells were washed twice and resuspended in FACS buffer until flow cytometric analysis.

Microglia for RNA-seq: Mice were terminally anaesthetized and perfused with ice-cold HBSS. Brain hemispheres were cut into small pieces with a razor blade and homogenized by Dounce in a solution of HBSS containing 15 mM HEPES, 0.5% glucose, 12.5 KU/ml DNase I, and 0.1 U/ml RNAase inhibitor. Brain homogenates were filtered through a 70-μm strainer followed by myelin removal using myelin removal beads (Miltenyi Biotec). Cells were resuspended in FACS buffer (2% FBS in PBS) and stained for 30 min with anti-CD45-PE (Thermo Fisher, clone 104) and anti-CD11b-Alexa Fluor 700 (Thermo Fisher, clone M1/70). Dead cells were excluded by staining with Live/dead Aqua (Thermo Fisher). Cell sorting was performed on an ARIA II (BD Biosciences) with FACSDiva software (BD Biosciences). $CD45^{low}CD11b^{+}$ cells were defined as the microglial population and were sorted directly into buffer RLT (Qiagen) and stored at -80 ˚C. The resulting microglia from two pooled mouse brains/sample yielded on average 80,000 $CD45^{low}CD11b^{+}$ cells and were processed for bulk RNA-seq experiments. FlowJo software was used for analysis and depiction of the gating strategy.

**Chemokine and cytokine multiplex assay**

Chemokine and cytokine analysis was carried out at the Human Immune Monitoring Core (Stanford University) or Eve Technologies (Calgary, Alberta, Canada) using magnetic bead-based multiplex Luminex assays (Cat# LXSAMSM, R&D Systems, Inc., Minneapolis, MN). Plates were read using a Luminex LabMap200 instrument. Mean fluorescence intensity (MFI) was averaged over duplicate wells for each cytokine per sample on each plate.

**Novel object recognition (NOR) task**

Novel object recognition was performed as previously described[13]. Briefly, mice were habituated to an empty arena (40 cm x 40 cm x 35 cm) containing wall-mounted visual cues for 5 minutes. Mice were then given one 5 min trial during which they explored two identical objects in fixed positions in the middle of the arena. Animals were randomly assigned to identical starting objects of either 25 mL cell culture flasks filled with sand or LEGO towers (3 cm x 3 cm x 7.5 cm). In the testing phase, mice explored the same arena for 5 min but with one object replaced by a novel, distinct object. Interactions with objects (sniffing or exploring within 2 cm of object; excluding time spent sitting on top of object) were manually timed in a blinded fashion to assess the fraction of time spent exploring each object (single object interaction time/total interaction time with both objects). Between trials, the arena and objects were thoroughly cleaned with 10% EtOH.

**Barnes maze task**

The Barnes maze test was performed as previously described with adaptations to accommodate aged and AD model mice[13]. Briefly, a large circular maze containing 16 holes on the outer edge was centered over a pedestal and elevated approximately 3 feet above the floor. All holes were open to the floor except for the escape hole. The escape hole consisted of a PVC elbow joint connector. Distinct visual cues were placed at four equally spaced points around the maze. An overhead light, two additional standing lights, and a fan blowing on the maze provided motivation to find the escape hole while also serving as supplemental visual cues. The escape hole position was fixed for all trials. Mice performed four trials per day for four consecutive days, as follows. For the adult WT, aged WT, and aged *Trem1-/-* cohorts, the starting location of the mouse was moved relative to the escape hole position every trial. For the 5XFAD and aged $APP^{Swe}$ cohorts, the starting location remained unchanged for all trials. Escape Latency is defined as the time for a mouse to enter the escape hole. If any mouse failed to locate the escape hole within 90 seconds, they were gently guided by light tapping towards the escape hole and assigned an escape latency score of 90 seconds. The maze and escape holes were thoroughly cleaned with 10% EtOH after every trial. Testing was performed by an investigator blinded to experimental conditions.

**Peritoneal Macrophages**

Peritoneal macrophages were collected from *Trem1-/-* and WT male mice. Mice were injected intraperitoneally with 1.5 ml 3% (w/v) thioglycolate medium (Cat# 211716, BD Biosciences) for Seahorse experiments, and peritoneal macrophages were isolated 3–4 days later by flushing with ice-cold 1× PBS buffer (Corning). For metabolomics and RNA-seq experiments, peritoneal macrophages were collected by lavage and enriched using CD11b MicroBeads (Cat# 130-049-601, Miltenyi Biotec, San Jose, CA) following the directions provided by the manufacturer. Cells were seeded at a density of $3 \times 10^6$ cells per well in DMEM supplemented with 10% heat-inactivated fetal bovine serum (FBS; Cat# F4235, Sigma-Aldrich), 100 U $ml^{-1}$ penicillin and streptomycin, and maintained at 5% $CO_2$ at 37 °C. After overnight culture, cells were washed twice with medium to remove nonadherent cells.

**RNA Sequencing and analysis**

Bulk RNA extraction and sequencing were performed by Novogene, Inc (Sacramento, CA). RNA degradation and contamination was monitored on 1% agarose gels and RNA purity was verified using the NanoPhotometer® spectrophotometer (IMPLEN, CA, USA). RNA integrity and quantification were assessed using the RNA Nano 6000 Assay Kit of the Bioanalyzer 2100 system (Agilent Technologies, CA, USA). Total mRNA was transcribed into full length cDNA using Takara V3 Stranded Prep kit (Cat.

634485, Takara Bio USA, Inc., Mountain View, CA). Clustering of index-coded samples was performed on a cBot Cluster Generation System using PE Cluster Kit cBot-HS (Illumina) according to the manufacturer's instructions. After cluster generation, the library preparations were sequenced on an Illumina platform and paired-end reads were generated. Raw data (raw reads) in FASTQ format were processed through fastp. In this step, clean data (clean reads) were obtained by removing reads containing adapter and poly-N sequences and reads with low quality. At the same time, Q20, Q30 and GC content of the clean data were calculated. All downstream analyses were based on clean data of high quality. Reference genome and gene model annotation files were downloaded from the genome website browser (NCBI/UCSC/Ensembl) directly. Paired-end clean reads were aligned to the reference genome using the Spliced Transcripts Alignment to a Reference (STAR) software (v2.6.1d). FeatureCounts (v1.5.0-p3) was used to count read numbers mapped for each gene. Differential expression analysis between two conditions/groups was performed using DESeq2 (v1.20.0) using default settings. DESeq2 provides statistical routines for determining differential expression in digital gene expression data using a model based on the negative binomial distribution. The resulting *P* values were adjusted using the Benjamini and Hochberg's approach for controlling the False Discovery Rate (FDR). Genes with an FDR-adjusted *P* value (*q*-value) < 0.05 by DESeq2 were designated differentially expressed. Enrichment analysis of DEGs was performed using the Gene Ontology (GO) enrichment analysis tools with the PANTHER classification system. Hypergeometric testing was performed to assess overrepresentation of DEGs ($q < 0.05$) within relevant gene sets including lysosome, phagocytosis, cytokines and chemokines.

**Seahorse Assay for Real-Time Oxygen Consumption Rate (OCR) and Extracellular Acidification Rate (ECAR)**

Mouse peritoneal macrophages or primary microglia were plated at $1.8 \times 10^5$ cells per well in a Seahorse XF24 Cell Culture Microplate (Agilent, Santa Clara, CA, USA). Cells were washed twice with Agilent Seahorse XF Media (Agilent) supplemented with 1 mM sodium pyruvate, 2 mM L-glutamine, and 10 mM D-glucose; a final volume of 500 µl was placed in each well. Cells were incubated in a 0% $CO_2$ chamber at 37°C for 1 hr before being placed into a Seahorse XFe24 Analyzer (Agilent). Cells were treated with 1 µM Oligomycin, 1 µM FCCP, and 0.5 µM rotenone/antimycin. A total of three OCR and pH measurements were taken per condition.

**Transmission Electron Microscopy (TEM)**

Primary peritoneal mouse macrophages ($2 \times 10^6$ cells/ml) were isolated. Briefly, cells were fixed in 2% glutaraldehyde and 4% paraformaldehyde in 0.1 M sodium cacodylate (pH 7.4), treated with 10% gelatin solution in sodium cacodylate buffer, and incubated with 2% osmium tetroxide. Cell pellets were stained with 1% uranyl acetate, dehydrated in ethanol, and embedded in resin (EPON epoxy resin). Sections were generated using a Leica EM UC7 ultramicrotome (Leica Microsystems) at 50 nm and placed within grids stained with a 1:1 mix of 3% uranyl acetate, 50% acetone for 30–60 s. Grids were imaged with a JEM 1400 transmission electron microscope (JEOL) at ×1,200 for low magnification and ×12,000 for high magnification (unless otherwise noted) using Gatan Microscopy Suite software (Gatan, Inc., Pleasanton, CA). Images were quantified by an individual blinded to experimental conditions in ImageJ. Abnormal mitochondria were defined as containing at least one of the following features: paracrystalline inclusions, linearization of cristae and abnormal angular features, concentric layering of cristae membranes, matrix compartmentalization, nanotunneling, in combination with doughnut-shaped or balloon-shaped mitochondria. Mitochondria measurements were completed using NIH ImageJ software (National Institutes of Health, USA) by investigators blind to experimental conditions.

**Targeted metabolomics**

Metabolites were extracted from CD11b-purified primary peritoneal macrophages in a 80:20 methanol:water solution in a volume of 75 mL solvent per 1-million cells, vortexed, incubated on dry ice for 10 min, and centrifuged at 16,000 g for 20 min, and the supernatant was assayed by LC-MS analysis. Extracts were analyzed within 24 hr by liquid chromatography coupled to a mass spectrometer (LC-MS). The LC–MS method involved hydrophilic interaction chromatography (HILIC) coupled to the Q Exactive PLUS mass spectrometer (Thermo Scientific)[17]. The LC separation was performed on an XBridge BEH Amide column (150 mm 3 2.1 mm, 2.5 mm particle size, Waters, Milford, MA). Solvent A was 95%: 5% H2O: acetonitrile with 20 mM ammonium bicarbonate, and solvent B was acetonitrile. The gradient was 0 min, 85% B; 2 min, 85% B; 3 min, 80% B; 5 min, 80% B; 6 min, 75% B; 7 min, 75% B; 8 min, 70% B; 9 min, 70% B; 10 min, 50% B; 12 min, 50% B; 13 min, 25% B; 16 min, 25% B; 18 min, 0% B; 23 min, 0% B; 24 min, 85% B; 30 min, 85% B. Other LC parameters are: flow rate 150 ml/min, column temperature 25°C, injection volume 10 mL and autosampler temperature was 5°C. The mass spectrometer was operated in both negative and positive ion mode for the detection of metabolites. Other MS parameters were: resolution of 140,000 at m/z 200, automatic gain control (AGC) target at 3e6, maximum injection time of 30 ms and scan range of m/z 75-1000. Raw LC/MS data were converted to mzXML format using the command line "msconvert" utility[18]. Data were obtained with MAVEN software[19,20].

Metabolomics data analysis was carried out using MetaboAnalyst version 5.0[21]. Metabolomic data were log-transformed and scaled according to the auto-scaling feature (mean-centered and divided by the standard deviation of each variable). Metabolites that were significantly different by ANOVA (with FDR correction) were subjected to hierarchical clustering analysis using the Euclidian distance measure and Ward clustering algorithm. Differentially expressed metabolites by volcano plot underwent pathway-based enrichment analysis using 84 metabolite sets based on KEGG human metabolic pathways in MetaboAnalyst 5.0.

**Integration of transcriptomic and metabolomic features**

Integration of macrophage RNA-seq and metabolomic data was performed using the Joint Pathway Analysis feature in MetaboAnalyst 5.0. Enrichment analysis was carried out by hypergeometric test with degree centrality selected as topology measure. A loose integration method was applied such that enrichment analysis was performed separately for genes and metabolites in their "individual universe" then the individual *P*-values were combined via weighted *Z*-tests. Weights were based on the overall proportion of each omics within the "universe". Significantly enriched pathways were determined by applying the cutoff of $q < 0.05$.

**Immunocytochemistry of NRF2 in peripheral macrophages**

Primary macrophages were collected by peritoneal lavage of ice-cold PBS. Cells were maintained in DMEM/F12 + 10% FBS + 1% PS solution at 5% $CO^2$ at 37°C. After overnight culture, cells were washed twice with media to remove nonadherent cells and were further cultured for an additional 24-hours. Cells were washed twice with PBS to remove media and cells were fixed with ice-cold 100% methanol for 12 minutes at -20°C. Cells were washed 3 x with PBS with 0.1% Tween 20, and then blocked for 1 hour with 1% BSA and 10% NGS with 0.1% Tween 20. Coverslips were incubated with antibodies to NRF2 (1:500, Cat# ab62352, Abcam) and CD11b (1:2000, Cat# NB600-1327, Novus Biologicals, Centennial CO). Coverslips were washed, stained with DAPI, and coverslipped as before. Z-stack images of CD11b+ cells spanning 15 µm were captured using Zeiss LSM 980 inverted confocal microscope using the 63x oil objective. In ImageJ, the DAPI channel was used to first construct a nuclear mask wherein NRF2 signal was measured within each z-plane.

**Primary Microglial Cultures**

10-16 P1-P3 C57/BL6 mice were euthanized and hippocampus and cortex were harvested, removing meninges under a dissection microscope. Brains were trypsinized for 25 min, with agitation every 5 min. Trypsinization was stopped with Trypsin inhibitor (0.6 mg/ml) in DMEM/F12 + 10% FBS + 1% PS solution. Tissue was homogenized and then passed through a 100 µm cell strainer. Cells were plated in a T-175 $cm^2$ flask and incubated in 5% $CO_2$, 37˚C incubator for 5 days in DMEM/F12 + 10% FBS + 1% PS. After 5 days, media was changed and cells were allowed to sit for another 9 days (14 days total, changing media every 5 days). Mixed cultures were then shaken for 8 h at 225 rpm and microglial cells were resuspended in

DMEM+10% FBS + 1% PS and plated at a confluency of 1.8 x $10^5$ per well in an XF24 Seahorse plate. Cells were treated with amyloid-beta oligomers (A$ß_{42}$ oligomers; 100 nM) or veh for 20 hr and Seahorse was carried out as described below.

**Immunofluorescent staining of mouse brain**

Coronal tissue sections containing the dorsal CA1 region of the hippocampus were washed with PBS containing 0.3% Triton X-100. Sections were blocked in 10% NDS for 1 hour then incubated overnight in primary antibody solution containing anti-Iba1 (1:1000, Cat# 019-19741, Wako) and 6E10 (1:1000, Cat# 803001, BioLegend, San Diego, CA) in 5% NGS. Sections were washed 3 x 10 min and incubated for 2 hours at RT with secondary antibodies (Abcam). Sections were again washed and stained with DAPI for 10 min. After a final wash in PBS, sections were mounted on glass slides and coverslipped using ProLong Gold Antifade reagent (ThermoFisher). For quantification of mature amyloid plaques, sections were incubated with Thioflavin S. Tissue was imaged using a Zeiss LSM 980 inverted confocal microscope running version 3.3 Zen Blue software (Zeiss, Dublin, CA). Images containing the CA1 region of the hippocampus were captured in z-plane spanning 15 µm with a step-size of 0.5 µm. An ImageJ macro was used to automatically z-project image stacks, despeckle, enhance contrast and skeletonize the IBA1 signal. The software plugin AnalyzeSkeleton (2D/3D) was used to calculate average and maximal branch length as previously described[14]. Representative images were prepared using ImageJ and Imaris software (Version 9.9.1, Oxford Instruments, Abingdon, UK).

For analysis of neuritic dystrophy, tissue sections spanning the dorsal hippocampus were washed 3x in PBS-T and then incubated for 20 minutes in 10 µMol X34 (SML1954, Milipore Sigma) solution prepared in 40% EtOH in PBS in 0.02N NaOH at room temperature. After incubation with X34, sections were washed in 40% EtOH and then PBS. Sections were then blocked for 1 hour in 10% NGS and incubated overnight in rabbit anti-BACE1 antibody (1:500, Cat#108394, Abcam). The following day tissue sections were washed and incubated with secondary antibody (Abcam). Sections were washed then mounted on glass slides and coverslipped with ProLong Gold antifade reagent (ThermoFisher). Slides were imaged using a LSM 980 Confocal microscope at 63X magnification. Three images were acquired each from 3-4 sections per mouse. Quantification of BACE1+ volume within 15 µm of X34+ amyloid plaque was done using Fiji (ImageJ) by experimenters blind to genotype information. Thresholded X34 images were binarized and ROIs were automatically selected according to the X34 plaque area. ROIs were expanded using the expand function to 15 µm. BACE1 levels in wild type mice were determined by applying a X34-expanded mask belonging to a 5XFAD mouse to the same hippocampal region. BACE1 was measured after thresholding within the ROIs using the Analyze Particles function and was recorded as % area.

**Synaptic mitochondrial isolation and Seahorse analysis**

Synaptic mitochondria were isolated as described previously[16]. Briefly, brain cortices were removed and added to cold freshly prepared 9 ml of mitochondrial isolation buffer (IB; 225 mM mannitol, 75 mM sucrose, 2 mM $K_2PO_4$, 0.1% BSA, 5 mM HEPES, 1 mM EGTA (pH 7.2)). Tissues were homogenized using a dounce homogenizer. The resultant homogenate was centrifuged at 13,000g at 4˚C and layered on top of 3 X 2-ml discontinuous gradient of 15%, 23% and 40% Percoll (GE) and centrifuged at 34,000g for 14 minutes at 4˚C. Following centrifugation, the band between 15% and 23% containing synaptosomes and band between 23% and 40% containing nonsynaptic mitochondria were removed and washed in IB with 0.02% digitonin (for synaptosomes only). The isolates were then pelleted by centrifugation at 16,500g for 15 minutes at 4˚C. The pellets were resuspended in IB and layered over another discontinuous gradient similar to that described above. Bands between 23% and 40% containing synaptic mitochondria were obtained and washed in ice cold IB. Protein estimation was performed using the Bradford assay (BioRad Laboratories). Isolated mitochondria were immediately used for Seahorse analysis. 10 ug of freshly isolated synaptic mitochondria were plated in XFe24 cell culture microplates in a volume of 50ul mitochondrial assay solution (MAS; 70 mM Sucrose, 220 mM mannitol, 10 mM $KH_2PO_4$, 5 mM $MgCl_2$, 2 mM HEPES, 1 mM EGTA and 0.2% BSA with 10 mM succinate and 2 uM rotenone) and attached to the wells by spinning down the plates at 2000 rpm at 4˚C. After attaching mitochondria to the plate wells, volume was brought up to 450 uL of MAS containing substrate. In the meantime, the Seahorse XF24 Flux Analyzer was equilibrated to 37˚C overnight a day before the assay. The final concentrations of substrates and inhibitors added to the wells were 4 mM ADP, 2.5 µg/ml Oligomycin A, 4 µM FCCP and 4 µM Antimycin A. The coupling assays were run in 4-5 replicate wells for each independent biological sample. XF24 data were collected according to Seahorse software (Agilent).

The basal OCR was determined in the presence of the incubation medium. The proton leak was determined after inhibition of mitochondrial ATP production by oligomycin, an inhibitor of the F0 F1 ATPase. The measurement of ATP production in the basal state was obtained from the decrease in respiration by inhibition of the ATP synthase with oligomycin. Then, the mitochondrial electron transport chain was stimulated maximally by the addition of the uncoupler FCCP. Finally, extra-mitochondrial respiration was estimated after the addition of the antimycin A and rotenone inhibitors of the complexes III and I, respectively. Coupling efficiency is the proportion of the oxygen consumed that drives ATP synthesis compared with that driving the proton leak and was calculated as the fraction of basal mitochondrial OCR used for ATP synthesis (ATP-linked OCR/basal OCR). Spare capacity is the capacity of the cell to respond to an energetic demand and was calculated as the difference between the maximal respiration and basal respiration.

**In vivo [$^{18}$F]FDG PET/CT imaging**

[$^{18}$F]FDG was obtained from the Cyclotron & Radiochemistry Facility at Stanford University. Quality control criteria were set and the tests were performed according to USP 823. In vivo [$^{18}$F]FDG PET/CT imaging was performed in 17-19 mo female mice. Mice were fasted overnight (12–14h) prior to imaging with access to water. Mice were anesthetized with isoflurane gas (2-3% in $O_2$) and blood samples acquired via tail prick were used to measure blood glucose levels in duplicates using a glucose meter (Accu-check) immediately prior to intraperitoneal injection with 292–322μCi of [$^{18}$F]FDG. Each mouse was anesthetized for a maximum of 5 minutes before being returned to home cage to allow tracer accumulation while awake. For PET/CT imaging, mice were anesthetized (3% for induction and 2% for maintenance in $O_2$) and 20 min static PET images were acquired at 75–95 min following [$^{18}$F]FDG injection using a dual microPET/CT scanner (Inveon, Siemens). A 3-dimensional ordered subsets expectation-maximum (3DOSEM, 2 iterations) and MAP-SP (18 iterations) reconstruction algorithm was applied to PET images (128 x 128 x 159 matrix size, 0.776 x 0.776 x 0.96mm voxel size). CT images were acquired to provide anatomical reference and to apply scatter and attenuation correction to PET data. PET images were analyzed using Vivoquant software (inviCRO) and visualized using Inveon Research Workspace (IRW, Siemens). Brain uptake was quantified as previously described using a brain atlas approach[15]. In brief, CT images were used to fit a 3-dimensional mouse brain atlas and to obtain [$^{18}$F]FDG uptake values in *a priori* regions of interest including the hippocampus and thalamus. PET uptake was expressed as SUVglc using the average blood glucose measurement for each mouse.

**Mendelian Randomization**

Two-sample Mendelian randomization (MR) was used to infer the causal effect of sTREM1 and sTREM2 on AD risk using the TwoSampleMR package[1,2] with summary statistics from genome-wide association studies (GWAS) testing association with SomaScan aptamers level in 35,559 Icelanders[3] and with AD in 75,024 cases and 397,844 controls[4]. MR inverse variance weighting was used to assess the significance of the association of the exposure with the outcome. MR Egger and MR Median Weighted estimators were used to confirm the robustness of these associations. Bidirectional MR was used to confirm the direction of causality.

**Quantitative immunoblotting of human AD brain**

Postmortem brain material was obtained from the Arizona Study of Aging and Neurodegenerative Disorders and Brain and Body Donation Program[5]. AD was defined as intermediate or high probability that dementia was due to AD according to NIA-Reagan criteria[6]. For immunoblot experiments, patients consisted of four groups stratified by Braak stage[7] including non-demented Braak I–II - zero to sparse plaques (n=12), non-AD demented Braak I–II - zero to sparse plaques (n=12), AD Braak III–IV- demented with moderate plaques (n=12), and AD Braak V–VI - demented with frequent plaques (n=12). We excluded cases with clinicopathologic evidence of Parkinson's disease (defined as having two of the three cardinal clinical signs of resting tremor, muscular rigidity and bradykinesia, along with pigmented neuron loss and Lewy bodies in the substantia nigra), dementia with Lewy bodies[8], progressive supranuclear palsy[9], motor neuron disease (including amyotrophic lateral sclerosis, primary lateral sclerosis and motor neuron disease associated with frontotemporal lobar degeneration), corticobasal degeneration (defined by the classic H&E histopathology of achromatic, swollen neurons in the cerebral cortex and indistinct inclusions within pigmented neurons of the substantia nigra as well as abnormal, phosphorylated tau or Gallyas-positive astrocytic plaques in the cerebral cortex), Pick's disease (defined as clinical dementia with tau or silver-stain positive Pick bodies within neurons of the cerebral cortex, hippocampus and/or basal ganglia), multiple system atrophy (defined by atrophy and gliosis of the cerebellar folia, basal pons, substantia nigra and/or striatum, as well as alpha-synuclein or silver-positive glial and neuronal cytoplasmic inclusions), and Huntington's disease (defined by the characteristic trinucleotide repeat expansion in the gene for huntingtin). Brains from subjects with cancer, sepsis, or stroke were also excluded. Post-mortem interval was less than 8 hours. Cases were balanced for sex in each group and all groups were matched for age at death. Demographic characteristics of the study participants are shown in **Supplementary Table 2**.

Frozen medial frontal gyrus (MFG) tissue was homogenized in 8x volume of lysis buffer (in 1% Triton X-100, 0.5% NP-40, 25 mM Tris HCl, 100 mM NaCl and protease and phosphatase inhibitors), sonicated, and centrifuged at 13,000 x g for 20 minutes. The protein concentration was determined using the BCA protein assay kit (Thermo Fisher Scientific, USA). Samples were separated on NuPAGE 4-12% Bis-Tris gels (Cat# WG1403A, Invitrogen) and transferred to polyvinylidene difluoride membranes (Cat# IPVH00010, Millipore Sigma, Burlington, MA). Membranes were blocked with Tris-buffered saline (TBS)/0.1% Tween-20/5% milk for 1 hour at room temperature (RT) and then incubated overnight at 4°C with primary antibodies against TREM1 (1:300; ab93717, Abcam, San Francisco, CA), β-Actin (1:10,000, A5441, Sigma-Aldrich, St. Louis, MO). Blots were washed 3 x 10 minutes with TBS-T and incubated for 2 hours with IRDye 800CW goat anti-rabbit (LI-COR Biosciences, Lincoln, NE) and IRDye 680and IRDye 680RD goat anti-mouse (LI-COR Biosciences) secondary antibodies (both 1:10,000). Blots were imaged using a LI-COR CLX-1306 instrument and analysis was completed using Image Studio Lite software (Version 5.2.5, LI-COR Biosciences). Human liver tissue lysate (Cat# HT-314, Zyagen, San Diego, CA) was used as positive control for human TREM1 and TREM2 proteins. Positive controls and full uncropped blots are presented in **Supplementary Fig. 8c-d**.

**Immunohistochemical (IHC) staining of human AD brain**

Formalin-fixed paraffin embedded (FFPE) frontal cortex from control and AD donors were obtained from University of Washington. Human healthy control and AD brain FFPE tissue sections were deparaffinized by heating at 56 °C for 1 hr and immediately passed through xylenes and a graded EtOH series from 100% I, 100% II, 95%, 70% into PBS (5 min incubation each step). Antigen retrieval was performed to reveal epitopes for antibody binding by incubating tissue in a 10 mM citrate buffer solution with 0.05% Tween 20 (pH 6.0) under low boil for 20 min. Sections were cooled to room temperature then washed with PBS. Endogenous peroxidase activity was quenched by incubating in 2% $H_2O_2$ solution for 20 min while shaking at RT. To further permeabilize the tissue, sections were incubated in two 10 min washes of PBS with 0.3% Triton-X shaking at RT. Tissue was blocked with 10% NDS (Jackson ImmunoResearch Laboratories, Inc.) in PBS for 1 hour shaking at RT and then incubated overnight at 4°C shaking in the following primary antibodies: rabbit polyclonal anti-TREM1 (Cat# 93717, Abcam) at 1:200 dilution and mouse monoclonal anti-β-Amyloid, 1-16 (clone 6E10, Cat# SIG-39320, Covance, Burlington, NC) at 1:1000 dilution. Tissue was then washed 3 x 10 min with PBS and incubated in secondary antibody solution for 2 hours while shaking at RT. Biotinylated goat anti-rabbit (1:1000, Cat# VA-1000, Vector Laboratories, Burlingame, CA) and biotinylated goat anti-mouse (1:1000, Cat# BA-9200, Vector Laboratories) were used as secondary antibodies. Sections were washed 3 x 10 min with PBS and then incubated for 1 hour with Avidin-Biotin complex (VECTASTAIN Elite, Cat# PK-6100, Vector Laboratories) shaking at RT. Sections were developed with the Vector 3,3'-diaminobenzidine (DAB) substrate kit (SK-4100). This process was repeated for the 6E10 antibody. The Vector VIP substrate kit (SK-4600) served as chromogen for the 6E10 antibody. Sections were dehydrated through a graded alcohol series, cleared in xylenes, and coverslipped with DPX mounting media (Cat# 44581, Sigma-Aldrich, St. Louis, MO). Photomicrographs were acquired using a Keyence BZ-X710 microscope running BZ-X Viewer Software (Keyence, Itasca, IL).

**Immunofluorescent staining of human brain**

Free-floating frontal cortex ribbons from Alzheimer's disease and control donors were obtained from the Arizona Study of Aging and Neurodegenerative Disorders and Brain and Body Donation Program. Antigen retrieval was performed for 1 hour at 37°C using 10 mM citrate buffer solution with 0.05% Tween 20 (pH 6.0). Sections were washed with PBS and then blocked in 10% normal donkey serum (NDS, Cat# 017-000-121, Jackson ImmunoResearch Laboratories, Inc., West Grove, PA) in PBS. Sections were then incubated overnight in rabbit polyclonal anti-TREM1 (Cat# 93717, Abcam, 1:150, 93717) and anti-CD163 at dilution of 1:25 in 5% NDS in PBS with 0.4% Tween. Tissue was washed and then incubated in secondary antibody solution (1:1000) in 2% NSD in PBS-T. Sections were incubated with 1% Thioflavin

S to identify amyloid plaques. Sections were washed, mounted on glass slides and coverslipped with Prolong Gold aqueous mounting media (Cat# P36930, ThermoFisher, Waltham, MA).

**Statistical analysis**

Means of two groups were compared using unpaired Student's *t*-tests. Pearson chi-square was used to compare differences in categorical variables. Normally distributed continuous variables were compared using one- and two-way ANOVA and Tukey's post hoc test. Non-normally distributed data were analyzed using nonparametric Kruskal-Wallis test followed by post hoc Mann-Whitney U test. All tests were two-sided. Human TREM1 and TREM2 levels were analyzed by ANCOVA with age and sex included in the model. Associations between TREM1 and TREM2 with pathological features were analyzed using linear regression with age and sex included as predictors. The statistical software packages used include R v.3.6.2 and Prism 8 and 9 (GraphPad Software). Data presented as mean ± SEM.

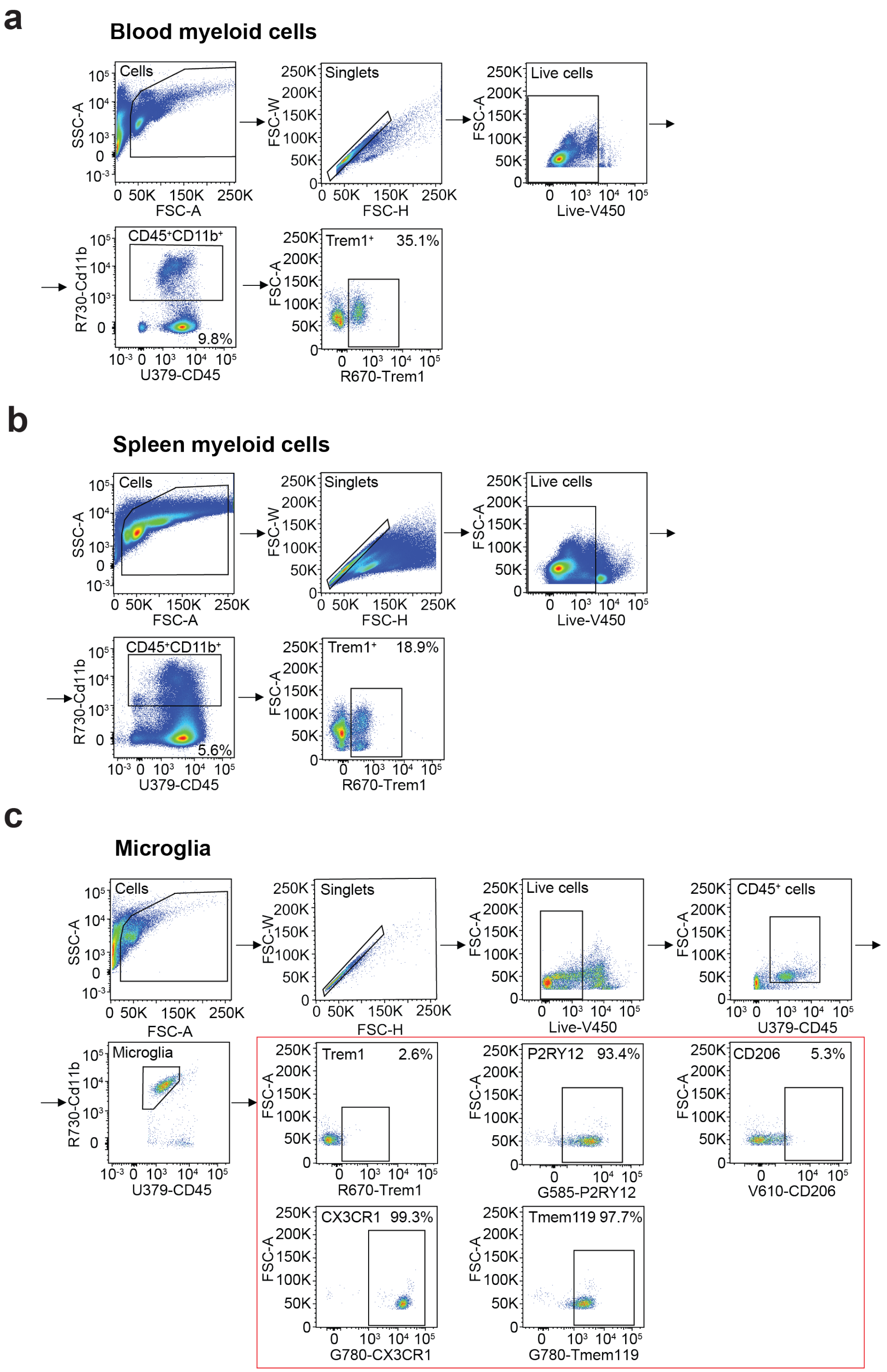


**Supplementary Fig. 1. Flow cytometry gating for quantification of TREM1 expression in blood, spleen and brain myeloid cells**

a. Gating strategy for live CD45$^+$Cd11b$^+$TREM1$^+$ blood myeloid cells
b. Gating strategy for live CD45$^+$Cd11b$^+$TREM1$^+$ spleen myeloid cells
c. Gating strategy for live CD45$^{lo}$Cd11b$^+$TREM1$^+$ microglia; also includes gating for P2RY12, CD206, CX3CR1 and Tmem119 microglial markers.

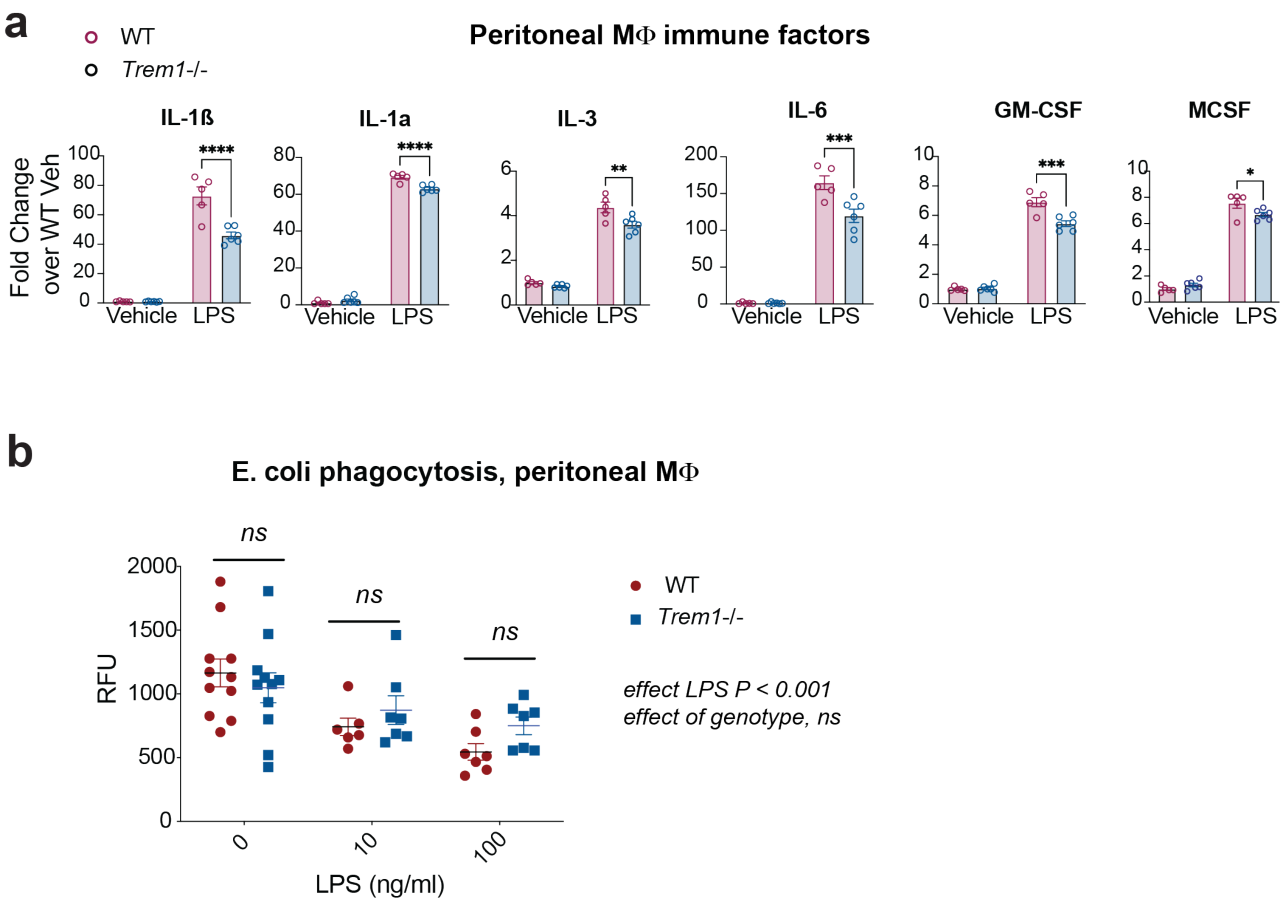


**Supplementary Fig. 2. TREM1 amplifies levels of inflammatory factors but does not alter phagocytosis**

a. Quantification of immune factors in lysates of peritoneal MΦ isolated from 8.5 mo WT and *Trem1*$^{-/-}$ mice treated with vehicle or 100 ng/ml LPS for 20 hr. Two-way ANOVA with Tukey's post hoc test, **P* < 0.05, ****P* < 0.001, *****P* < 0.01 (n=5-6 male mice per genotype).

b. *Trem1* deficiency does not alter phagocytosis in peritoneal MΦ. Cells were stimulated with vehicle or *E.coli* +/- LPS 100 ng/ml for 20h. 2-way ANOVA, effect of LPS ***P < 0.001, no effect of *Trem1* genotype basally or with LPS stimulation. 2-3 mo male mice.

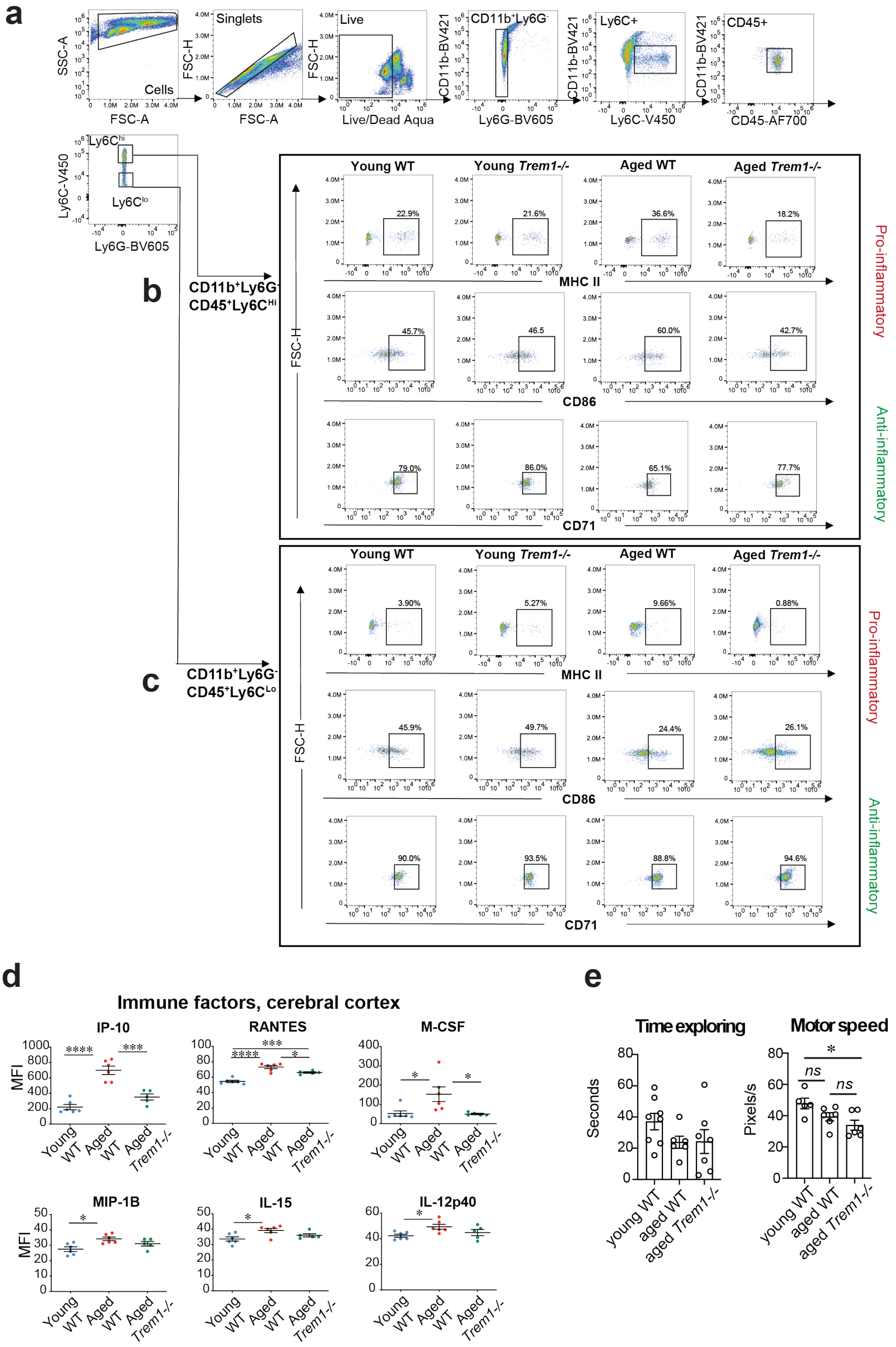


**Supplementary Fig. 3. TREM1 promotes a pro-inflammatory polarization state in aged peritoneal MΦ**

a. Gating strategy for live $Cd11b^+Ly6G^-CD45^+Ly6C^{Hi}$ and $Cd11b^+Ly6G^-CD45^+Ly6C^{Lo}$ peritoneal MΦ from young WT and *Trem1*$^{-/-}$ (2 mo) and aged WT and *Trem1*$^{-/-}$ (22-23 mo) male mice.
b. Representative gating for MHC II, CD86, and CD71 positive $Ly6C^{Hi}$ peritoneal MΦ.
c. Representative gating for MHC II, CD86, and CD71 positive $Ly6C^{Lo}$ peritoneal MΦ.
d. Multianalyte Luminex quantification of immune factors in young WT (2 mo) and aged WT (23-24 mo) and aged *Trem1*$^{-/-}$ (23-24 mo) cerebral cortex. ANOVA with Tukey's post hoc test; $*P < 0.05$, $***P < 0.001$, $****P < 0.01$ (n=5-6 male and female mice per group).
e. (Left) Total time exploring (in seconds) during the training phase of the NOR task; ANOVA with Tukey's post-hoc test is not significant. (Right) Speed (pixels/second) of mice during the final trial of the Barnes Maze; ANOVA with Tukey's post hoc test; $*P < 0.05$, (n=5-6 male and female mice per group).

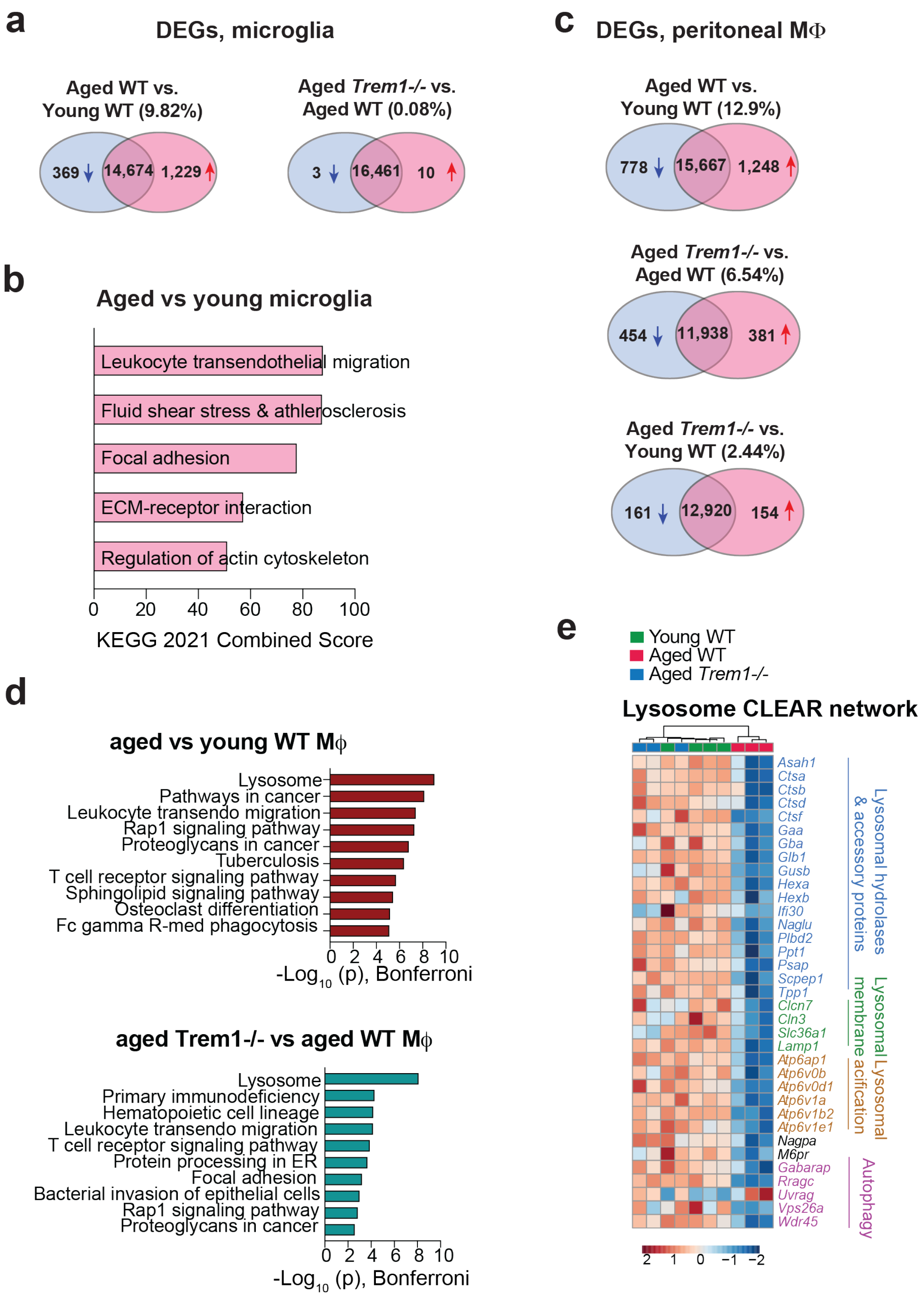


**Supplementary Fig. 4. *Trem1* deficiency alters gene expression in aged peripheral MΦ with minimal changes in aged brain microglia**

a. Venn diagram of DEGs (all genes $q < 0.05$) in pairwise comparisons of microglia isolated from aged (18 - 20 mo) vs young (3 mo) mice and pairwise comparisons of microglia isolated from aged *Trem1*$^{-/-}$ vs aged WT mice (18-20 mo). Blue indicates number of downregulated genes while red indicates the number of upregulated genes (n=3 microglial samples (2 pooled male mice per sample) per group).

b. KEGG Pathway enrichment analysis of upregulated genes in aged compared to young microglia; there were no enriched pathways in the comparison of aged WT vs aged *Trem1*$^{-/-}$ microglia.

c. Venn diagrams showing the number of DEGs (all genes $q$ < 0.05) in pairwise comparisons of primary mouse MΦ from young WT (2 mo), aged WT (25 mo) and aged *Trem1-/-* (25 mo) mice. Blue indicates number of downregulated genes while red indicates the number of upregulated genes (n=3-4 male mice per group).

d. Pathway enrichment analysis of upregulated genes in aged MΦ vs young and in aged *Trem1*$^{-/-}$ vs aged WT MΦ.

e. Heatmap of DEGs ($q$ < 0.05) comprising the Coordinated Lysosomal Expression and Regulation (CLEAR) network. Lysosomal functional categories are indicated for each gene. Aged *Trem1*$^{-/-}$ mice cluster with the young WT mice. Scale represents z-score values from FPKM. n=3-4 male mice per group.

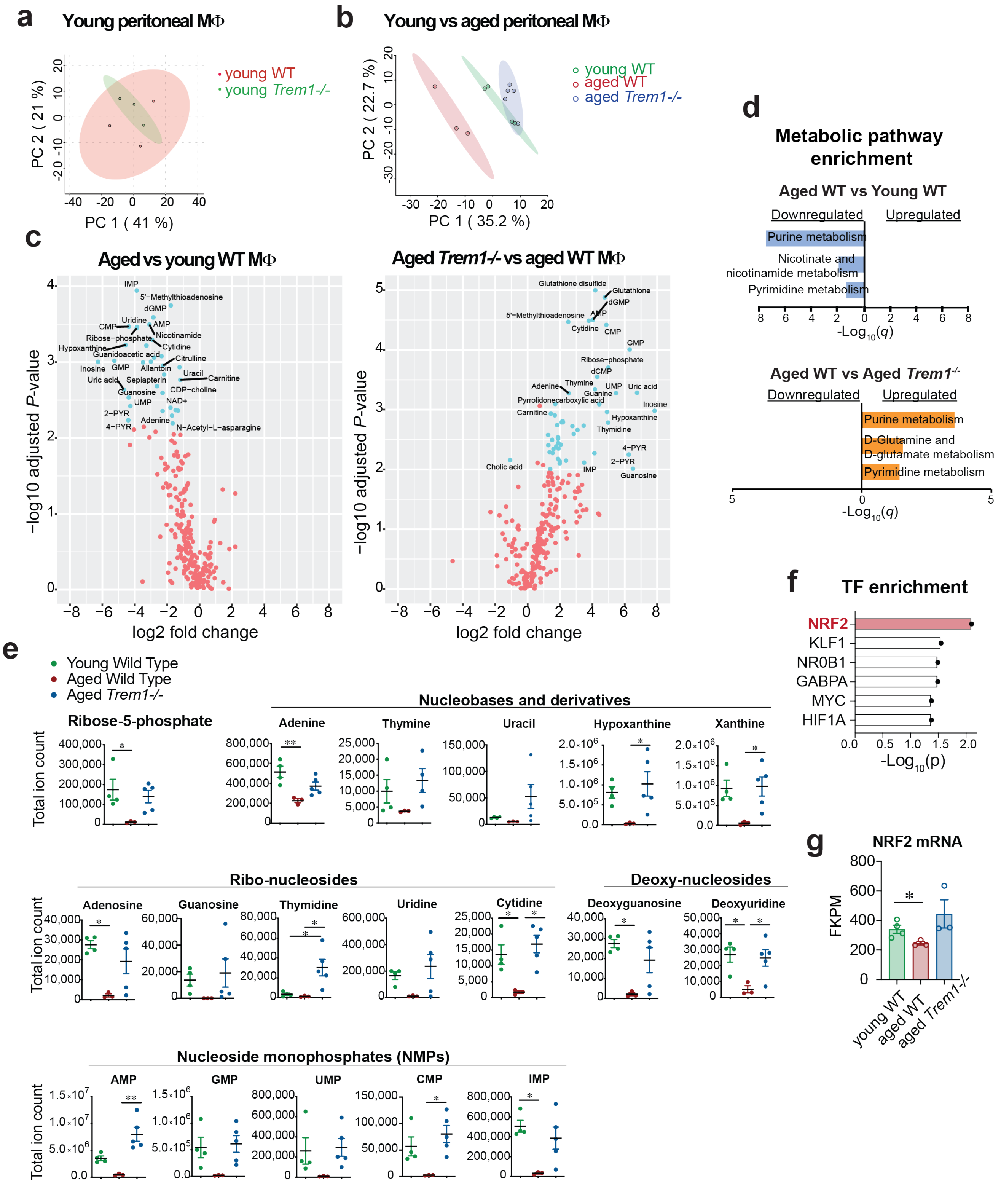


**Supplementary Fig. 5. TREM1 deficiency restores youthful metabolism in aged macrophages**

**a.** PCA of metabolites from young (2 mo) WT and *Trem1*$^{-/-}$ peritoneal MΦ (n=3 male mice per group).

**b.** PCA of significantly regulated metabolites from primary peritoneal MΦ isolated from young WT (2 mo), aged WT (25 mo) and aged *Trem1*$^{-/-}$ (25 mo) male mice. n=3-5 male mice per group.

**c.** Volcano plot comparing aged WT vs young WT (left) and aged *Trem1*$^{-/-}$ vs aged WT (right) peritoneal MΦ with Log2 fold change (FC) and -Log10(*P*) values of metabolites. Significantly regulated metabolites with $-\log_{10}(P) > 2$ and $\log_2$ fold change > 1 are shown in blue.

**d.** KEGG metabolic pathway enrichment analysis of differentially regulated metabolites between aged WT and young WT macrophages and between aged *Trem1*$^{-/-}$ vs aged WT MΦ. X-axis shows -Log10(*q*) value. No pathways were significant following FDR correction for the comparison of aged *Trem1*$^{-/-}$ vs young WT. Pathway analysis was performed using MetaboAnalyst 5.0; n=3-5 male mice per group.

**e.** Total ion count of Ribose-5P, the precursor of purines and pyrimidines and total ion counts for nucleobases and derivatives, ribo-nucleosides, deoxynucleosides, and nucleoside monophosphates. Data are analyzed by one-way ANOVA with Tukey's post hoc test. **P* < 0.05, ***P* < 0.01 (n=3-5 male mice per group). Abbreviations: AMP: adenosine monophosphate, GMP: guanosine monophosphate, UMP: uridine monophosphate, CMP cytidine monophosphate, IMP: inosine monophosphate.

**f.** Transcription factor (TF) enrichment analysis revealing TFs enriched for differentially expressed metabolite enzymes.

**g.** FKPM for NRF2. ANOVA followed by Tukey's post hoc test, **P* < 0.05 (n=3-4 male mice per group).

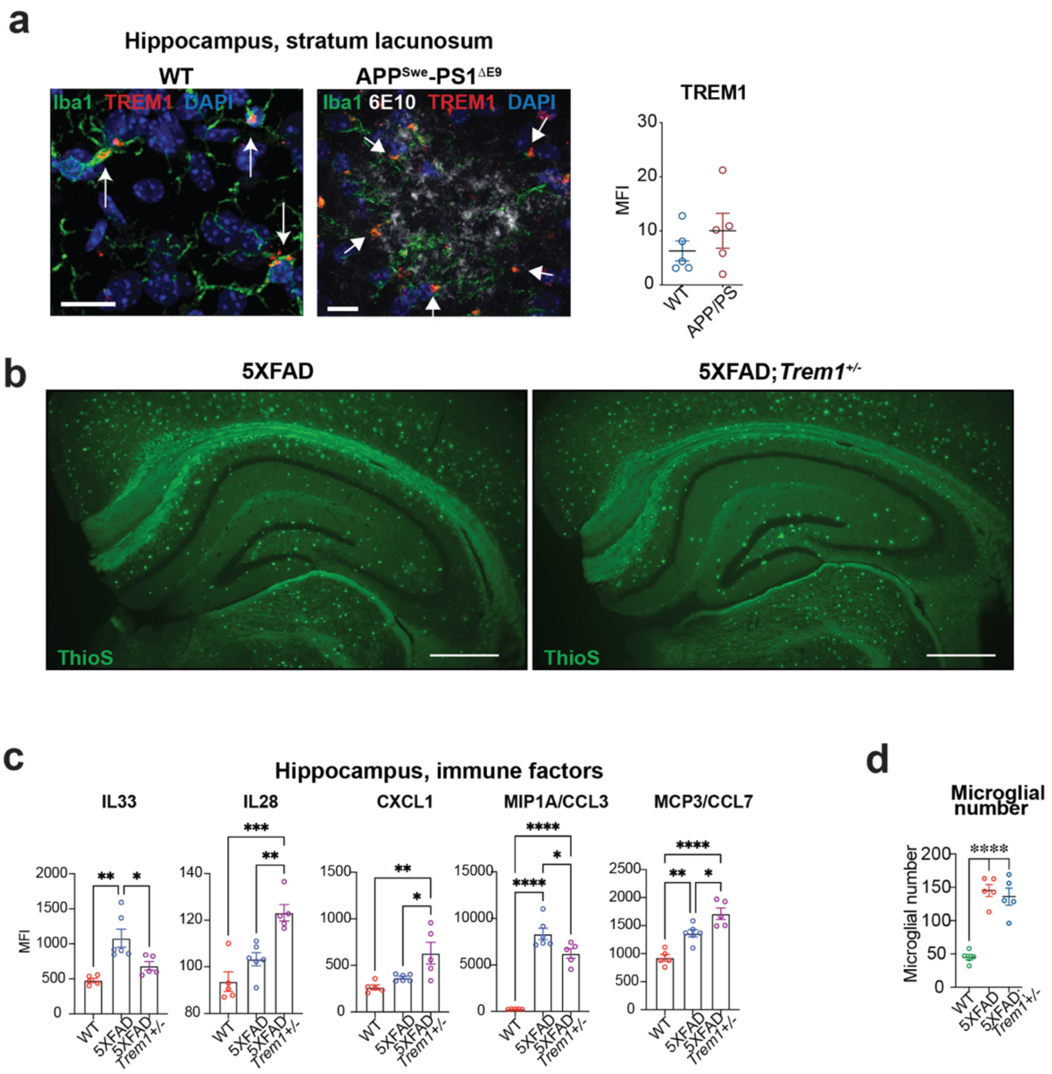


**Supplementary Fig. 6. TREM1 expression does not significantly change in context of accumulating amyloid.**

- **a.** (Left) Immunofluorescent staining of TREM1 in 9 mo WT and *APP*$^{Swe}$-*PS1*$^{\Delta E9}$ hippocampal stratum lacunosum in IBA1+microglia. 5-7 hippocampal sections per mouse were imaged, n=5 mice/genotype. Scale bar = 10 µm. TREM1 colocalizes with Iba1+ microglia (white arrows). (Right) Quantification of MFI shows no significant differences between genotypes.
- **b.** Thioflavin-S (ThioS) fluorescent staining of hippocampus in 9-10 mo 5XFAD and 5XFAD;*Trem1*$^{+/-}$ female mice. Scale bar = 500 µm.
- **c.** Quantification of hippocampal immune factors using multianalyte Luminex in WT, 5XFAD, and 5XFAD;*Trem1*$^{+/-}$ in 10 mo mice. One-way ANOVA with Tukey post-hoc comparisons, *$P < 0.05$, **$P < 0.01$, ***$P < 0.001$, ****$P < 0.01$ (n= 5-6 female mice per group).
- d. Microglial number from Fig. **4i**. One-way ANOVA with Tukey's post hoc test. **** $P < 0.0001$ (n=5 male mice per condition).

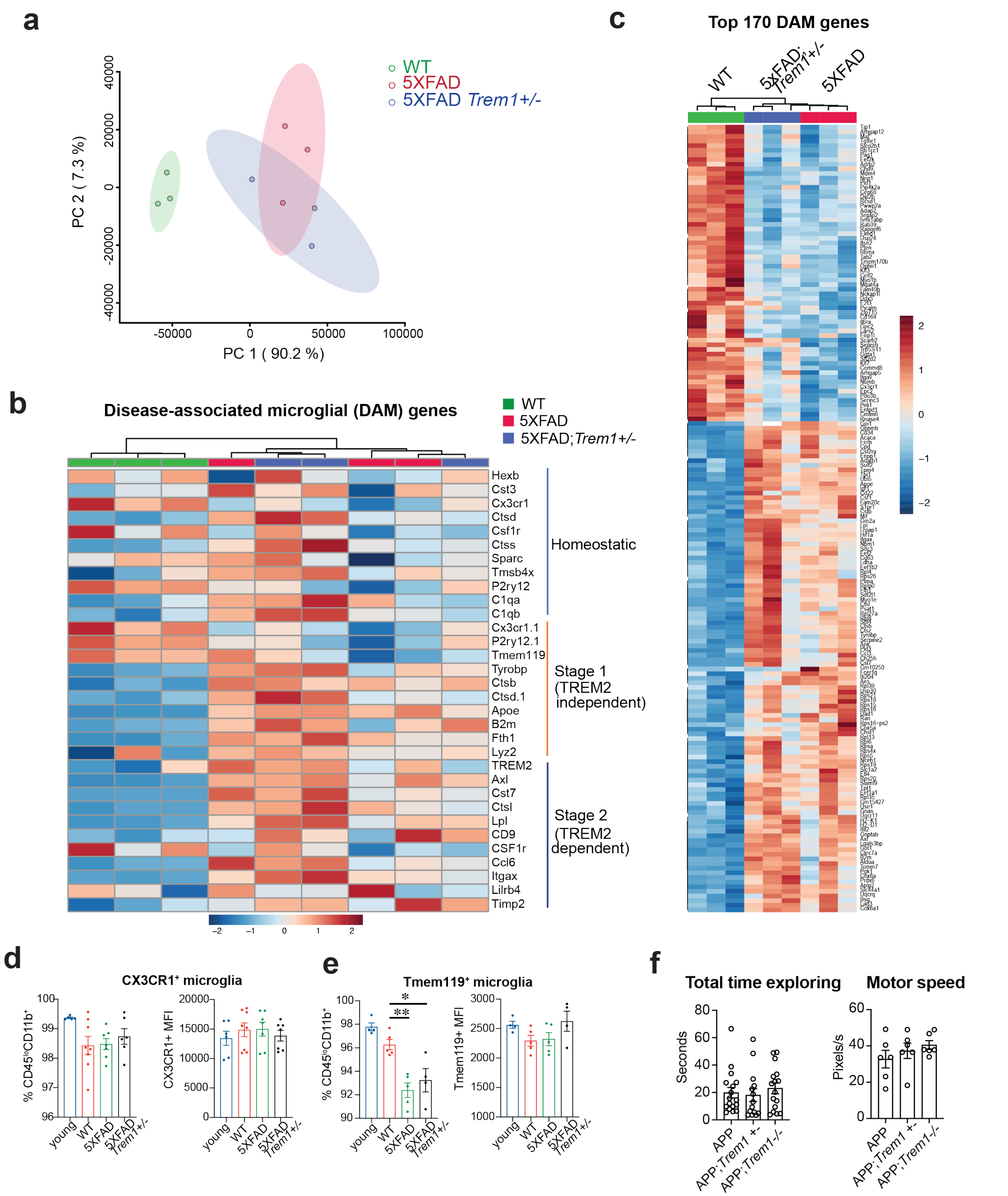


**Supplementary Fig. 7. Disease-associated microglial (DAM) signature in 5XFAD;*Trem1*$^{+/-}$ mice**

**a.** PCA of significantly regulated DAM signature genes from primary microglia isolated from WT, 5XFAD and 5XFAD/*Trem1*$^{+/-}$ mice. Individual microglial samples were pooled from 2 male mice (n=3 samples per condition).

**b.** Hierarchical clustering of DAM signature gene set showing homeostatic, DAM Stage 1 and DAM Stage 2 genes. Scale represents z-score values from FPKM.

**c.** Hierarchical clustering of top 170 DEGs (FDR-corrected) belonging to the full DAM signature gene set. Scale represents z-score values from FPKM. Scale represents z-score values from FPKM.

**d.** (Left) Percent of CX3CR1+ microglia in young (3 mo) and 13-17 mo WT, 5xFAD and 5xFAD;*Trem1*$^{+/-}$ mice. (Right) MFI of CX3CR1+ microglia. One-way ANOVA with Tukey's post hoc test, non-significant (n= 5-8 male and female mice per group).

**e.** (Left) Percent of Tmem119+ microglia in young (3 mo) and 13-17 mo WT, 5xFAD and 5xFAD;*Trem1*$^{+/-}$ mice. (Right) MFI of Tmem119+ microglia. One-way ANOVA with Tukey's post hoc test, *$P < 0.05$, **$P < 0.01$ (n=4-5 mice per group).

**f.** (Left) Total time exploring (in seconds) during the training phase of the NOR task in *APP*$^{Swe}$ mice; ANOVA with Tukey's post-hoc test (n=14-17 male and female mice per condition). (Right) Motor speed (pixels/sec) during the final training session of the Barnes maze; ANOVA with Tukey's post-hoc test (n=6 male and female mice per group).

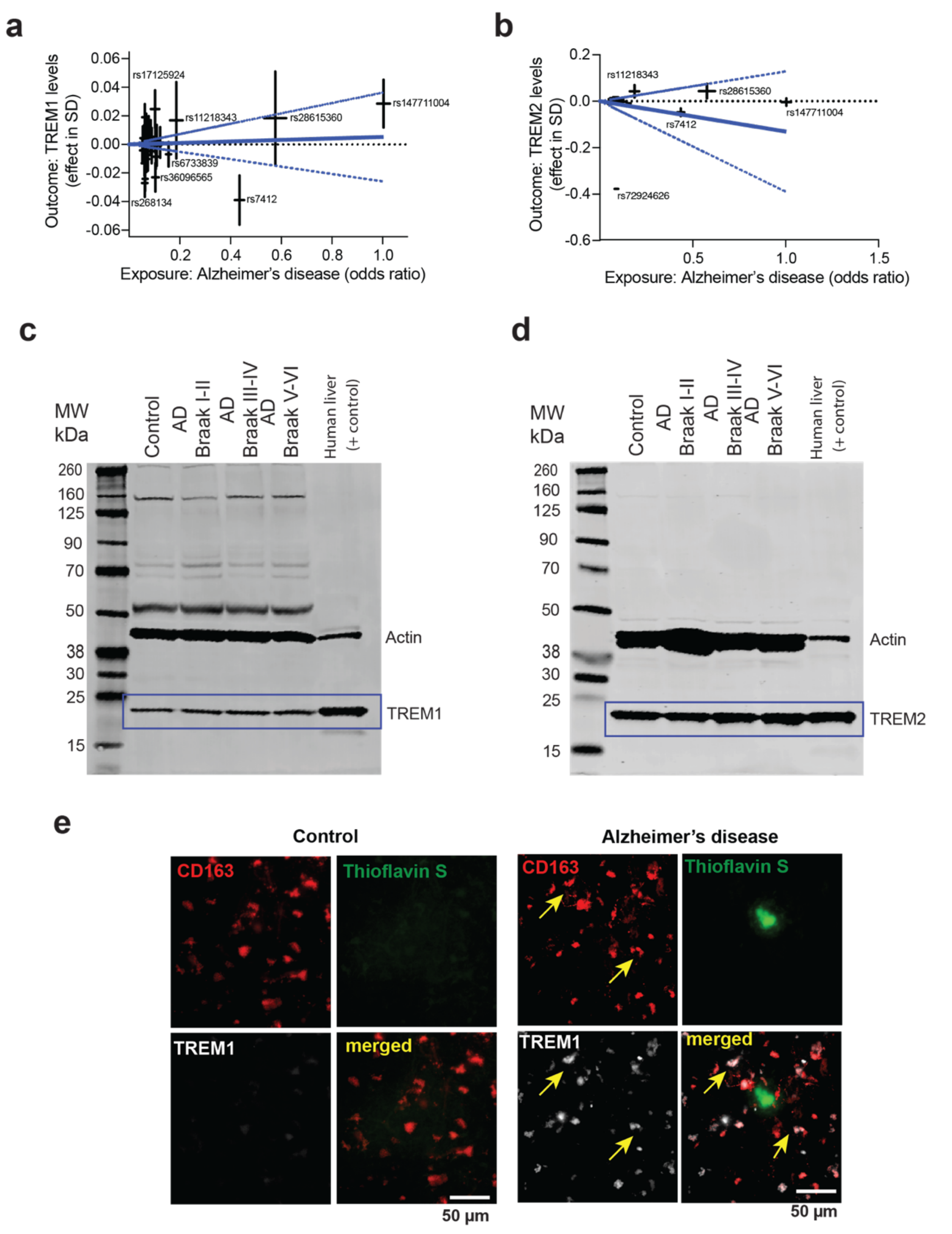


**Supplementary Fig. 8. TREM1 expression in development of AD**

**a.** Mendelian Randomization (MR) analyses with Alzheimer's disease (AD) risk level as exposure and sTREM1 plasma protein level as outcome. Blue lines are estimated MR-median weight effects, and the dashed lines indicate the 95% confidence interval for the MR effects.

**b.** Mendelian Randomization (MR) analyses with AD risk level as exposure and TREM2 plasma protein levels as outcome. Blue lines are estimated MR-median weight effects, and the dashed lines indicate the 95% confidence interval for the MR effects.

**c.** Immunoblot of TREM1 protein in postmortem human mid frontal gyrus. Clinicopathological diagnoses: non-demented Braak I-II, demented non-AD Braak stages I-II, AD Braak III-IV and AD Braak V-VI (n=12 donors per group). Primary antibody detected human TREM1 band at the molecular weight of positive control (human liver lysates, Cat# HT-314, Zyagen, San Diego, CA). The TREM1 band used for analysis are indicated by blue box. Also shown is the band for ß-actin at 42 kDa.

**d.** Immunoblot of TREM2 protein in postmortem human mid frontal gyrus. Primary antibody detects human TREM2 band at the molecular weight of positive control (human liver lysates, Cat# HT-314, Zyagen, San Diego, CA). TREM2 bands used for analysis are indicated by blue box. Also shown is the band for ß-actin at 42kDa.

**e.** Immunofluorescent colocalization of CD163, a marker of perivascular macrophages (red), Thioflavin S (green) and TREM1 (white) in frontal cortex free floating sections from control and AD brain. Diagonal yellow arrows indicate colocalization of CD163 and TREM1 staining. Scale bar = 50 μm.

**Supplementary Table 1: List of Metabolites Detected by LC-MS**

| Metabolite |
| --- |
| 5-Methoxytryptamine |
| 1-Methyl-Histidine |
| 1-Methyladenosine |
| 2-Furoylglycine |
| 2-Phenylbutyric acid |
| 2-PY |
| 2-PYR |
| 3-Hydroxyanthranilic acid |
| 3-Methyl-Histidine |
| 4-Guanidinobutanoic acid |
| 4-PY |
| 4-PYR |
| 4-Trimethylammoniobutanal |
| 5-Methoxytryptophan |
| 5-Phenylvaleric acid |
| 5'-Methylthioadenosine |
| acetyl-CoA |
| adenine |
| adenosine |
| ADP |
| Aminoadipic acid |
| AMP |
| Anserine |
| anthranilate |
| arginine |
| aspartate |
| ATP |
| carnitine |
| Carnosine |
| CDP |
| CDP-choline |
| citrulline |
| creatine |
| Creatine phosphate |
| cytidine |
| D-Fructose-6-phosphate |
| D-glucosamine-1/6-phosphate |
| deoxyadenosine |
| deoxyguanosine |
| deoxyuridine |
| dGDP |
| dGMP |
| DL-2-Aminooctanoic acid |
| FAD |
| GDP |
| glucosamine-6-phosphate |
| glucose-1-phosphate |
| glutamate |
| glutamine |
| GMP |
| GSH:GSSG |
| guanine |
| histidine |
| hydroxyproline |
| IDP |
| Imidazoleacetic acid |
| IMP |
| Indole-3-ethanol |
| inosine |
| kynurenic acid |
| kynurenine |
| L-Aspartyl-L-phenylalanine |
| Linoleic acid |
| lysine |
| m-Coumaric acid |
| Melatonin |
| methionine |
| Methionine sulfoxide |
| N-Acetyl-L-alanine |
| N-Acetyl-L-arginine |
| N-Acetyl-L-asparagine |
| N-acetyl-L-glutamine |
| N-Acetyl-L-isoleucine |
| N-Acetyl-L-leucine |
| N-Acetyl-L-proline |
| N-Acetylcarnitine |
| N-Acetyllysine |
| N-Acetylserine |
| N-Acetylserotonin |
| N-Octanoylglycine |
| NAD+ |
| NADP+ |
| NADPH |
| NaR |
| 1-Methyl imidozolacetic acid hydrochloride |
| 2-Amino-3-phosphonopropionic acid |
| 2-Furoylglycine |

2-Hydroxybutyric acid
2-Methyl-3-ketovaleric acid
2-Phenylbutyric acid
2-PY
3-Cresotinic acid
3-Hydroxyanthranilic acid
3-hydroxylbutyrate
3-Methyladipic acid
4-PY
5-Aminopentanoic acid
5-Hydroxymethyluracil
5-Phenylvaleric acid
7-methylguanosine
a-ketoglutarate
acetoacetate
acetylphosphate
aconitate
Aconitic acid
Adonitol
ADP
ADP-D-glucose
alanine
allantoin
Alloisoleucine
Aminoadipic acid
Aminolevulinic acid
Anserine
anthranilate
Arachidic acid
arginine
asparagine
aspartate
ATP
b-alanine
b-Aminoisobutyric acid
betaine
Carnosine
CDP
Cholic acid
citrate/isocitrate
CMP
creatine
Creatine phosphate
Creatinine
D-2-Aminobutyric acid
D-erythrose-4-phosphate
D-Fructose-6-phosphate
D-Phenyllactic acid
D-Xylose
dCMP
deoxyribose-phosphate
dGDP
dimethylglycine
Dimethylmalonic acid
DL-Valine
FAD
Fructose
fructose-1-6-bisphosphate
fumarate
Galactose
Galacturonic acid
GDP
Glucose
glucose-1-phosphate
glutamate
glutamine
Glutaric acid
glutathione
glutathione disulfide
Glyceraldehyde-3-phosphate
glycerate
glycine
glycolate
Glycyl-L-leucine
Glycyl-L-proline
Guanidoacetic acid
guanine
guanosine
histidine
Homocarnosine
homoserine
Hydroxyphenyllactic acid
hydroxyphenylpyruvate
hydroxyproline
hypoxanthine
IDP
IMP
isoleucine
L-2-Hydroxyglutaric acid
lactate
Lactose
leucine
Linoleic acid

lysine
m-Coumaric acid
malate
Malonic acid
Mandelic acid
methionine
Methionine sulfoxide
Methylglutaric acid
Methylmalonic acid
Methyloxovaleric acid (Ketoleucine)
Methylsuccinic acid
myo-inositol
Myristic acid
N-Acetyl-L-alanineneg
N-Acetyl-L-asparagine
N-Acetyl-L-isoleucine
N-Acetyl-L-leucine
N-Acetylaspartic acid
N-Acetylcarnitine
N-Acetylglycine
N-Methyl-D-aspartic acid
N-Octanoylglycine
NAD+
nicotinamide
nicotinate
O-acetyl-L-serine
O-Phosphoethanolamine
ornithine
orotate
p-Cresol
pantothenate

Parahydroxyphenylacetic acid
phenylalanine
phosphoenolpyruvate
Pimelic acid
proline
Pyroglutamic acid
Pyrrolidonecarboxylic acid
pyruvate
quinolinate
Retinal
Retinoic Acid
sarcosine
Sepiapterin
serine
shikimate
sn-glycerol-3-phosphate
Stearic acid
Suberic acid
succinate
Taurodeoxycholic acid
threonine
thymine
Tiglic acid
trans-Glutaconic acid
tryptophan
tyrosine
UDP
UDP-D-glucose
Undecanoic acid
uracil
Uric acid

uridine
Urocanic acid
valine
xanthine
nicotinamide
nicotinamide mononucleotide
nicotinamide riboside-2
O-acetyl-L-serine
O-Phosphoethanolamine
ornithine
pantothenate
phenylalanine
Pipecolic acid
Pyroglutamic acid
Pyrrolidonecarboxylic acid
Retinal
Retinoic Acid
ribose-phosphate
Ribothymidine
taurine
Taurodeoxycholic acid
threonine
thymidine
thymine
Tryptamine
tryptophan
UDP
UMP
uridine
Xanthurenic acid

### Supplementary Table 2: Demographic Characteristics of Study Participants

| Variable | Total (*n* = 48) | Non-Demented Braak I-II (*n*=12) | Demented Braak I-II (*n*=12) | AD Braak III-IV (*n*=12) | AD Braak V-VI (*n*=12) | *P* |
|---|---|---|---|---|---|---|
| Gender (F/M), no. (%) | 23 (48)/25 (52) | 5 (42)/7 (58) | 6 (50)/6 (50) | 6 (50)/6 (50) | 6 (50)/6 (50) | ns, a |
| Age, years (mean +/- SD) | 83.2 (4.1) | 81.8 (4.1) | 82.4 (4.0) | 83.2 (4.3) | 84.3 (3.9) | ns, b |
| PMI (hours) | 3.14 (1.14) | 2.95 (0.27) | 3.38 (0.48) | 3.24 (0.32) | 2.99 (0.21) | ns, b |
| Braak Stage (median, +/- SD) | 2.5 (1.71) | 1.5 (0.52) | 2 (0.29) | 4 (0.58) $ | 6 (0.51) & | *P*<0.0001, c |
| CERAD Neuritic Plaque Density (median, +/- SD) | 2.5 (0.99) | 1 (0.83) | 2 (0.45) @ | 3 (0.45) & | 3 (0.00) & | *P*<0.0001, c |

a Chi-Square test

b One Way ANOVA

c Kruskal-Wallis test

#, *P*<0.01 vs Non-Demented Braak I-II

$, *P*<0.001 vs Non-Demented Braak I-II

&, *P*<0.0001 vs Non-Demented Braak I-II
